# Expansion of RNA sequence diversity and RNA editing rates throughout human cortical development

**DOI:** 10.1101/2021.06.10.447947

**Authors:** Ryn Cuddleston, Laura Sloofman, Lindsay Liang, Enrico Mossotto, Xuanjia Fan, Minghui Wang, Bin Zhang, Jiebiao Wang, Nenad Sestan, Bernie Devlin, Kathryn Roeder, Joseph D. Buxbaum, Stephan J. Sanders, Michael S. Breen

## Abstract

Post-transcriptional modifications by RNA editing are essential for neurodevelopment, yet their developmental and regulatory features remain poorly resolved. We constructed a full temporal view of base-specific RNA editing in the developing human cortex, from early progenitors through fully mature cells found in the adult brain. Developmental regulation of RNA editing is characterized by an increase in editing rates for more than 10,000 selective editing sites, shifting between mid-fetal development and infancy, and a massive expansion of RNA hyper-editing sites that amass in the cortex through postnatal development into advanced age. These sites occur disproportionally in 3’UTRs of essential neurodevelopmental genes. These profiles are preserved in non-human primate and murine models, illustrating evolutionary conserved regulation of RNA editing in mammalian cortical development. RNA editing levels are commonly genetically regulated (editing quantitative trait loci, edQTLs) consistently across development or predominantly during prenatal or postnatal periods. Both consistent and temporal-predominant edQTLs co-localize with risk loci associated with neurological traits and disorders, including attention deficit hyperactivity disorder, schizophrenia, and sleep disorders. These findings expand the repertoire of highly regulated RNA editing sites in the brain and provide insights of how epitranscriptional sequence diversity by RNA editing contributes to neurodevelopment.

## MAIN

Over the course of development, DNA-encoded genes are precisely transcribed into functional RNA sequences, which undergo a variety co- and post-transcriptional modifications. Post-transcriptional modifications by RNA editing are a major contributor to the global diversity of RNA sequences^1,2^ and are predicted to occur at over a hundred million sites in the human transcriptome^3-5^, with most impacting neuronal genes^6,7^. While RNA editing is essential for neurodevelopment, there is little understanding of the temporal dynamics of RNA editing during human brain development. For instance, it remains unclear to what degree the frequencies and rates of RNA editing sites are developmentally regulated and when and where these sites are established from early fetal through late postnatal periods. Moreover, the genetic regulation of RNA editing through brain development periods also remains poorly resolved.

Adenosine-to-Inosine (A-to-I) editing is the most prevalent form of RNA editing in the brain, whereby inosine is recognized by the cellular machinery as guanosine during translation^8^. A-to-I editing can occur either at a single isolated adenosine, termed “selective editing”, or across many neighboring adenosines within an extended region or cluster, termed “RNA hyper-editing”^9-11^, and both are driven by adenosine deaminases acting on RNA (ADAR) enzymes. The vast majority of RNA editing occurs in primate-specific Alu elements^1,10^. These base-specific changes recode amino acid sequences of proteins, alter the locations of start or stop codons^12^, influence neuronal splicing patterns^13^, affect the ability of miRNAs to bind to their target sites and can alter the stability of the RNA secondary structures^14,15^. A requisite for neuronal development and function, several sites are known to drive essential physiological processes, including: tight regulation of Ca^2+^ permeability (*GRIA2*, Q/R)^16,17^, acceleration of recovery rates from desensitization (*GRIA2-4*, R/G)^18^, actin cytoskeletal remodeling at excitatory synapses (*CYFIP2*, K/E)^19,20^, gating kinetics of inhibitory receptors (*GABBAR*, I/M)^21,22^, maintenance of action potential for voltage-dependent potassium channels (Kv1.1, I/V), and calmodulin binding mediating inhibitory Ca^2+^-feedback on channels (Cav1.3).

Recognizing their importance, the developmental properties of recoding sites have been extensively examined in murine models of neurodevelopment and report a precise increase in editing levels from embryonic to postnatal periods^23-27^. It has been proposed that such editing activity is not solely explained by ADAR expression, but by the presence of a strict regulatory network that controls editing during development. Fewer studies have profiled editing activity across human brain development, and leverage either a small number of samples, a narrow developmental window, or study a select number of editing sites^28-32^. Markedly, one report uncovered 742 RNA editing sites that increased in editing levels during postnatal periods^31^ and a separate investigation also found an increasing pattern of global editing activity across *in vitro* maturation of hiPSC-derived brain organoids^32^; offering a partial view of RNA editing throughout neurodevelopment. Still, the developmental and regulatory features of RNA editing in human brain have not been fully compiled. Such information is essential to understand the biological roles of RNA editing throughout the widely different neurodevelopmental stages, from early differentiating cellular populations to the broad range of structures found in the adult brain.

To address these gaps, we present a systematic evaluation of both selective RNA editing and RNA hyper-editing in the developing human cortex. Our study covers a wide-range of developmentally distinct samples (**Figure 1**). We anchor our analysis around BrainVar data^33^, which comprises paired whole-genome and bulk tissue RNA-sequencing (RNA-seq) data from the dorsolateral prefrontal cortex (DLPFC) of 176 donors spanning prenatal and postnatal developmental periods, from six post-conception weeks (PCWs) to young adulthood (20 years). The frontal cerebral wall was assayed in nine brains before ten PCWs. These data were integrated with *in vitro* human embryonic and induced pluripotent stem cell models of corticogenesis and hundreds of cortical samples from advanced ages. We identify a dramatic shift in selective RNA editing and hyper-editing between mid-fetal development and infancy, which refines the timing of these events and the degree to which each editing site is involved. Our analytic framework illustrates evolutionary conservation of these profiles in the non-human primate and murine models of neurodevelopment and elucidates interactions between RNA editing sites and genes essential for neuronal differentiation, mRNA metabolism, synaptic signaling, receptor sequences and those important for neurodevelopment. We also uncover the genetic regulation of RNA editing and identify hundreds of *cis*-editing quantitative trait loci (edQTLs) and classify their effects as prenatal-predominant, postnatal-predominant, or constant across brain development. Several edQTLs co-localize with risk variants for neuropsychiatric and neurodevelopmental disorders. Data and analyses we present here refine our understanding of the RNA editing-mediated mechanism throughout cortical development and will provide significant benefits to the fields of neuroscience, genomics, and medicine.

**Figure 1.**
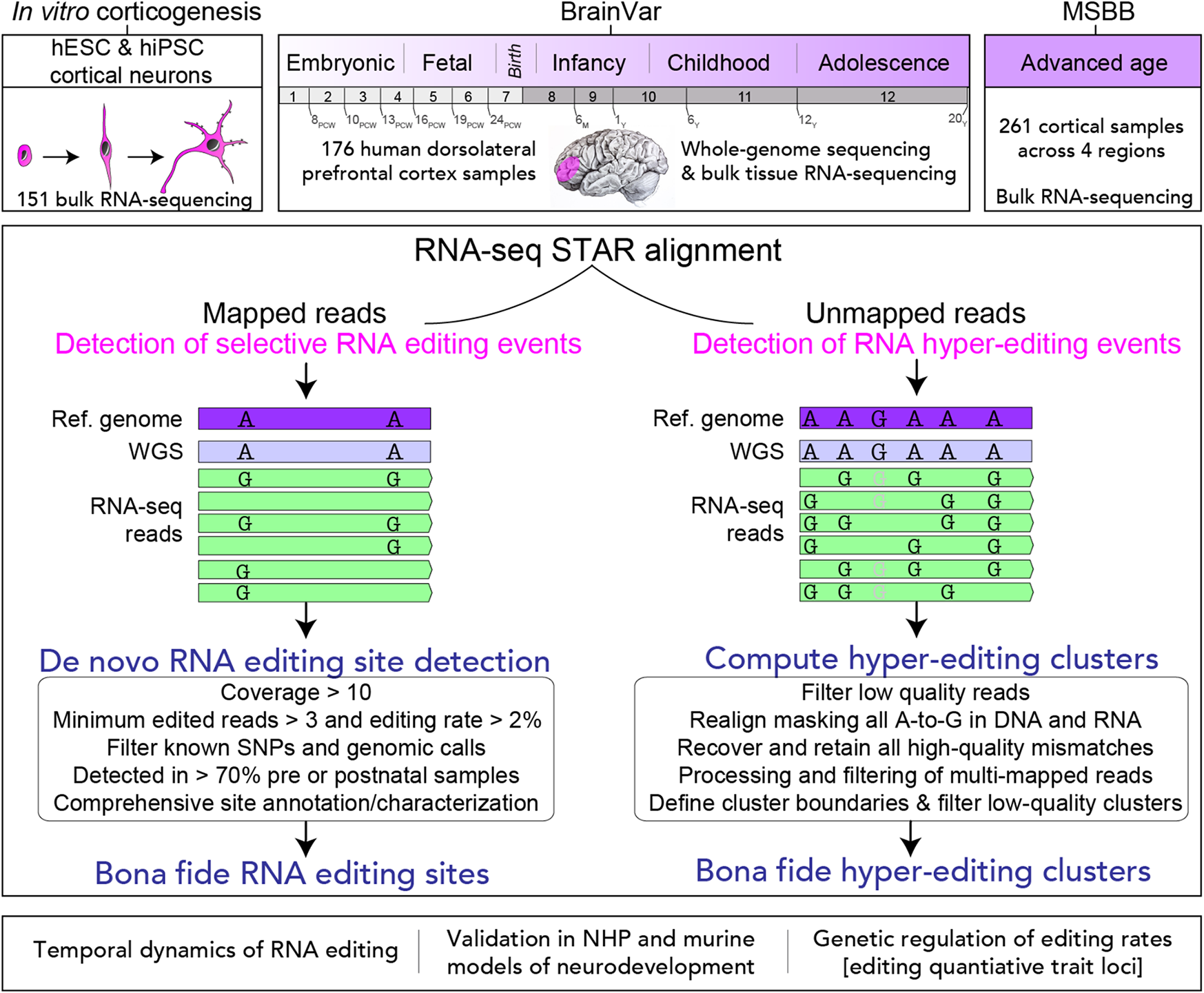
A general overview of the current study. Our analysis is anchored around paired WGS and RNA-seq data from BrainVar (*n*=176, DFLPC), together with *in vitro* human embryonic stem cell (hESC, *n*=24) and human pluripotent stem cell (hiPSC, *n*=127) models of corticogenesis and cortical samples from advanced stages of aging (*n*=261, 61-108 years). We also provide validation in a smaller developmental bulk brain transcriptome dataset from the forebrain (*n*=55) and hindbrain (*n*=59). We evaluate both selective RNA editing events from mapped RNA-seq bam files and RNA hyper-editing events from unmapped RNA-seq fastq files (*see Methods*). Our analytical approach sought to identify developmental trends in editing rates, provide evolutionary context in animal models of neurodevelopment, and elucidate common genetic variants that exert their effects by altering editing levels.

### Increased rate of selective editing through human cortical development

We computed an Alu editing index (AEI) as a measure of global selective A-to-G editing activity for each individual defined as the total number of selective A-to-G mismatches over the total coverage of adenosines in Alu elements (**Supplemental Table 1, Figure S1**). A threefold increase in global selective A-to-G editing rates occurred during cortical development (*p*=2.2×10^−28^, Cohen’s d=2.55), with a major shift between the mid-fetal period and infancy (**Figure 2A**). In parallel, *ADAR1* (*p*=2.9×10^−12^), *ADAR2* (*p*=3.1×10^−5^), and *ADAR3* (*p*=0.009) showed strong postnatal bias in expression (**Figure 2B**). Cellular deconvolution of bulk RNA-seq profiles revealed a decrease in the proportions of immature/fetal neurons and an increase in mature neurons from prenatal to postnatal periods (**Figure 2C**). Variation in the AEI was largely explained by age (*R*^*2*^=4.2%), followed by proportions of mature neurons within a sample (*R*^*2*^=3.3%), *ADAR2* (*R*^*2*^=0.49%) and *ADAR1* expression (*R*^*2*^=0.44%) (**Figure S2**). To provide deeper cell-specific information, we turned to single-nuclei RNA-seq of the DLPFC covering mid-fetal and postnatal periods. These data confirm ADAR expression levels are postnatally biased and that both *ADAR1* and *ADAR2* are highly expressed in mature excitatory neurons, whereas *ADAR3* shows a strong preference for a small subclass of inhibitory neurons (**Figure S3**). In keeping with the estimates of AEI variation, a significant postnatal bias in global editing activity remained after correcting for the proportion of mature neurons (*p*=2.1×10^−6^).

**Figure 2.**
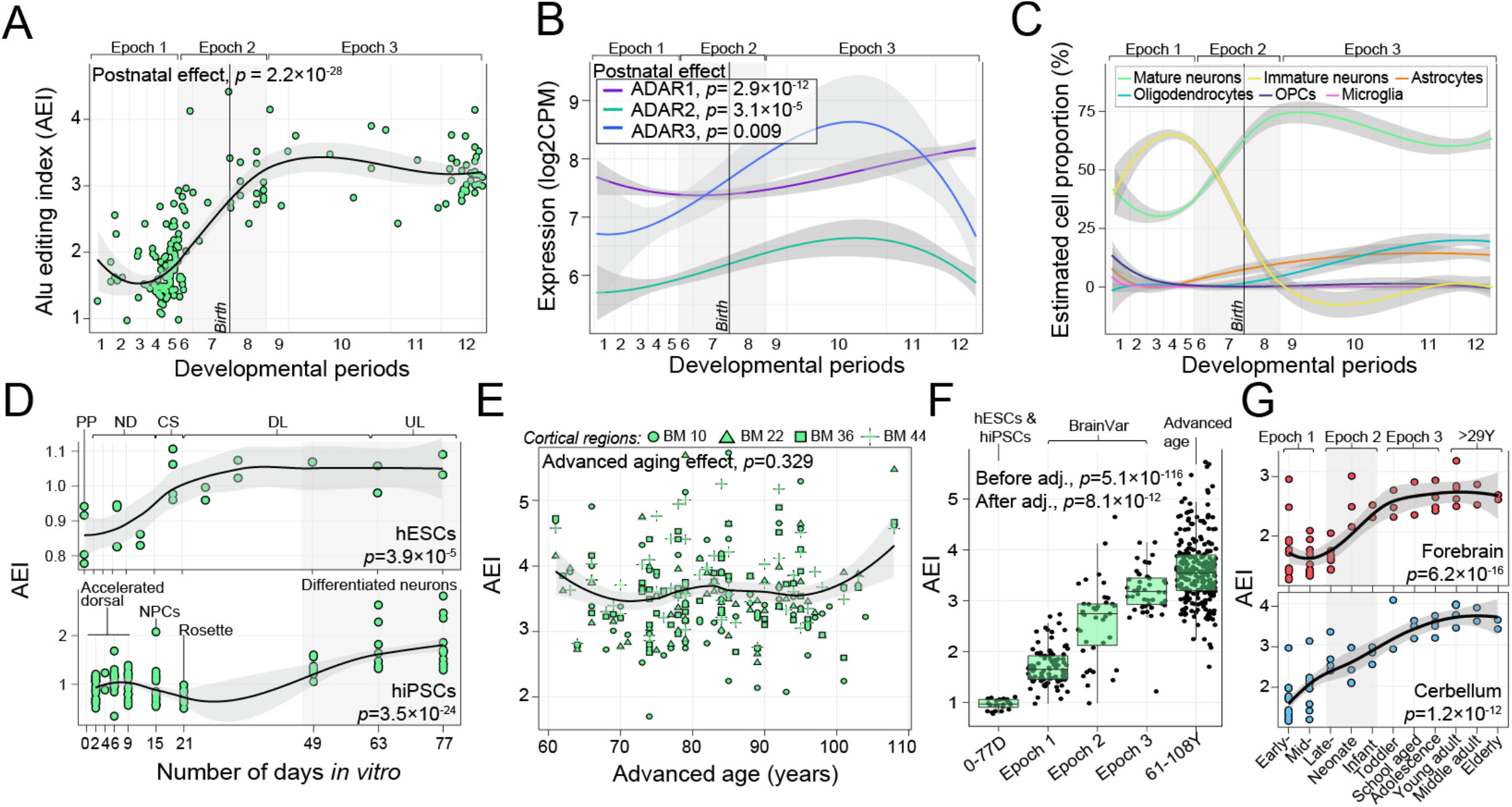
Alu editing index through cortical development. (**A**) The Alu editing index (AEI; y-axis), (**B**) expression profiles for *ADAR1, ADAR2* and *ADAR3* (y-axis) and (**C**) estimated cell type proportions based on cell-specific signatures from Darmanis et al., 2015 (y-axis) were computed for each DLPFC sample in BrainVar and examined across 12 developmental periods (log age, x-axis). Periods 1-7 reflect prenatal windows and periods 8-12 reflect postnatal windows. A linear regression was used to test for significance. (**D**) The AEI (y-axis) in human embryonic stem cell (hESC) and human induced pluripotent stem cell (hiPSC) models of corticogenesis across 77 *in vitro* (x-axis). (**E**) The AEI (y-axis) across advanced ages in the MSBB cohort (61-108 years). (x-axis). (**H**) The AEI (y-axis) across early hESCs, BrainVar (1^st^, 2^nd^ and 3^rd^ Epochs), and into advanced aging (MSBB). A linear regression was used to test for significance both with and without adjustment for neuronal cell type proportions. **(G**) Validation of the AEI in independent bulk forebrain (*n*=55) and cerebellar (*n*=59) bulk tissue transcriptome data across the lifespan.

We juxtaposed global selective A-to-G editing rates from BrainVar prenatal periods (Epoch 1) with two *in vitro* models of corticogenesis: 1) a human embryonic stem cell (hESC) model, spanning early stages of pluripotency (day 0) through upper layer generation (day 77); and 2) a human induced pluripotent stem cell (hiPSC) model, including 127 samples covering early differentiating cells (days 2-9) through mature differentiated neurons (day 77) (**Supplemental Table 1**). Both hESCs (*p*=3.9×10^−5^, Cohen’s d=1.44) and hiPSCs (*p*=3.5×10^−24^, Cohen’s d=1.67) captured this increase in global selective A-to-G editing (**Figure 2D**), and these rates were comparable across the two models. Notably, editing rates during late differentiation stages (days 49-77) in hiPSCs were similar to those in fetal brain tissue (Epoch 1) of BrainVar (*p*=0.38) (**Figure S4**).

Next, global selective A-to-G editing rates from BrainVar postnatal periods (Epoch 3) were contrasted with 261 cortical samples from older adults (61-108 years) (**Figure S5, Supplemental Table 1**). While the AEI was not dynamically regulated throughout advanced aging (*p*=0.39) (**Figure 2E**), editing activity steadily increased compared to Epoch 3 of BrainVar (*p*=3.1×10^−5^) (**Figure 2F**), suggesting that cortical global editing rates in the cortex might peak between 20-59 years of life. To test this, we explored the AEI in a smaller independent dataset of bulk forebrain (*n*=55) and cerebellar (*n*=59) tissues spanning fetal and postnatal periods (4 PCWs to 58 years) (**Supplemental Table 1**). Both the bulk forebrain and cerebellar tissues confirmed the increased rate of editing throughout development (*p*=6.2×10^−16^, Cohen’s d=2.97; *p*=1.2×10^−12^, Cohen’s d=2.44, respectively) and peak during young adulthood (∼30 years) (**Figure 2G**). We also show that editing activity in the brain displays the strongest developmental increase relative to four other human tissues sampled across the lifespan (**Figure S6**). Overall, global editing activity rises throughout maturation from early progenitors through fully differentiated neurons in the adult brain, even after adjustment for neuronal proportions (**Figure 2G**).

### Systematic characterization of selective editing events

To catalogue high-confidence selective RNA editing sites, a series of detection-based thresholds were imposed and all common and private genomic variants were removed (*see Methods*) (**Figure S7**). Using the resulting high-quality RNA editing sites and a sliding window spanning three neighboring developmental periods, a uniform number of sites per developmental window was observed (**Figure 3A**). RNA editing sites within each window were then compared to all remaining windows and a convergence of sites consistently edited either during prenatal or postnatal developmental periods was identified (**Figure 3B**). Based on this result, we aggregated RNA editing sites across i) prenatal samples only, ii) postnatal samples only, and iii) all available samples. We identified 3,879 prenatal predominant sites, 6,969 postnatal predominant sites and 10,652 ‘common’ sites with high detection rates across all cortical samples (**Figure 3C, Supplemental Table 2**). The majority of postnatal predominant and common sites were edited with ∼25% editing efficiently, while editing rates of prenatal predominant sites were significantly higher (**Figure 3D**).

**Figure 3.**
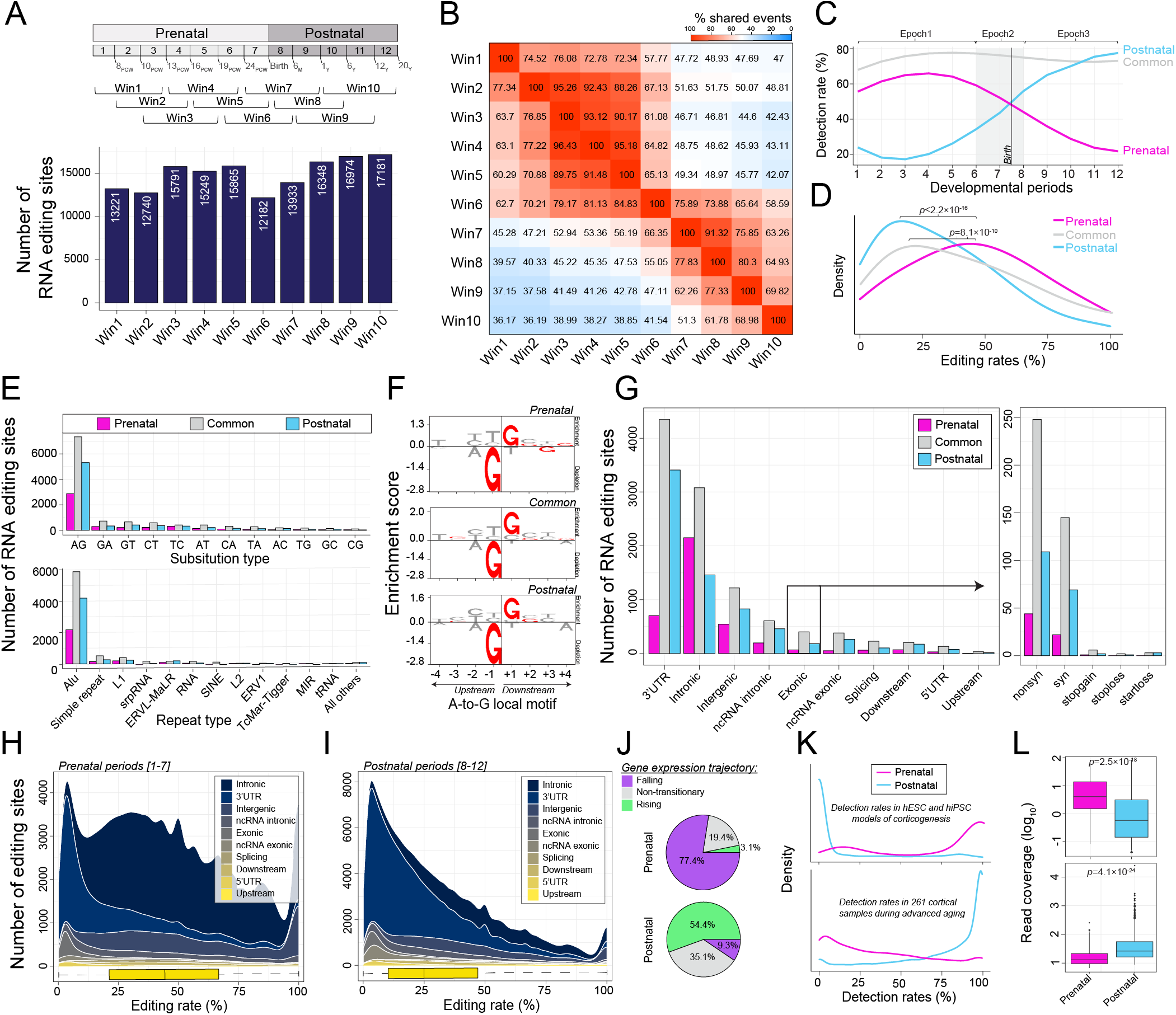
Identification and annotation of selective RNA editing sites. (**A**) A sliding window approach identified sites shared across 80% of samples per window, and a uniform distribution of editing sites were detected across all windows. (**B**) Sites specific to each window where then overlapped onto each other to reveal an enrichment of prenatal-specific (N sites=3,879) and postnatal-specific (N sites=6,969) sites, as well as sites found across all periods, termed ‘common sites’ (N sites =10,652). (**C**) Detection rates (%) indicate the fraction of samples for which prenatal, postnatal and common sites are detected with sufficient coverage per period. (**D**) Distribtuion of RNA editing rates (%, x-axis) across all prenatal, postnatal and common sites. Significance was computed using a Mann-Whitney U test. (**E**) Detected sites were annotated according to substitution type (upper) and to repeat elements (lower). (**F**) A-to-G sites enriched (y-axis) for a conserved local sequence motif, featuring a depletion of guanosine +1bp upstream and enrichment of guanosine -1bp downstream the target adenosine. (**G**) Sites according to genic region and an enrichment of 3’UTR sites among postnatal-specific sites. Distribution of editing rates among (**H**) prenatal-specific and (**I**) postnatal-specific sites by genic region. Here the amount of area reflects a larger contribution of editing sites from the corresponding genic region. (**J**) Gene expression trajectories for prenatal-specific sites largely decrease through development and trajectories for postnatal-specific sites largely increase through development. (**K**) Detection rates (%, x-axis) of prenatal- and postnatal-specific sites in *in vitro* models of corticogenesis (upper panel) and 261 cortical samples of advanced age (lower panel) illustrate a high detection rate (**L**) and high read coverage (log_10_ scale) of these corresponding temporal editing sites. Mann-Whitney U test was used to test for differences in coverage rates.

Validating the accuracy of our approach, these sites showed consistent hallmark signatures of RNA editing: i) The vast majority were A-to-G editing events (69% prenatal, 89% postnatal), mapped to Alu repeats (71.2%) and were known sites catalogued in editing databases (70.3%) (**Figure 3E**); ii) We confirmed the local sequence motif for all A-to-G editing events in which guanosine was depleted -1bp upstream and enriched +1 downstream the targeted adenosine (**Figure 3F**); iii) The majority of sites occurred in 3’UTRs and introns with few sites in coding regions (**Figure 3G-I**), with a significant enrichment of 3’UTR editing during postnatal periods (*p*=2.7×10^−6^) (**Figure S8**); iv) Most prenatal predominant sites mapped to genes with a falling expression trajectory over development (77.4%), while postnatal predominant sites mapped to genes with a rising expression (54.4%) (**Figure 3J**); v) We confirmed the temporal specificity for ∼45% prenatal predominant sites (*n*^*sites*^=1,745) in hESC and hiPSC models of corticogenesis (which were absent during advanced age) and ∼89% of postnatal predominant sites (*n*^*sites*^=6,132) in advanced age (which were absent hESCs and hiPSCs) (**Figure 3K-L**).

### Developmental regulation of selective editing events

For the 10,562 *bona fide* selective RNA editing sites in common across all BrainVar samples, differences associated with developmental age had the largest genome-wide effect on editing rates, followed by ADAR expression and neuronal proportions (**Figure S9**). Principal components accurately distinguished prenatal from postnatal samples based on editing rates for these sites (**Figure 4A**). Differential editing identified developmentally-regulated sites: 6,450 showed higher editing postnatally (‘postnatal biased’), 776 prenatally (‘prenatal biased’), and 3,426 were constant (‘unbiased’) across cortical development (**Figure 4B, Supplemental Table 3**). While these results were robust to adjusting for mature neuronal cell proportions, adjusting for *ADAR1* and *ADAR2* resulted in twofold reduction of developmentally regulated sites (**Figure S9, Supplemental Table 3**). Notably, the majority of postnatally biased RNA editing sites (∼86%) were known, while the majority of prenatally biased RNA editing sites (∼81%) were novel, despite similar sample numbers and the same thresholds for detection (**Figure 4C**). Novel sites occurred across all genic regions in a distribution similar to known sites and showed significantly more supporting read coverage compared to known sites (**Figure S10**).

**Figure 4.**
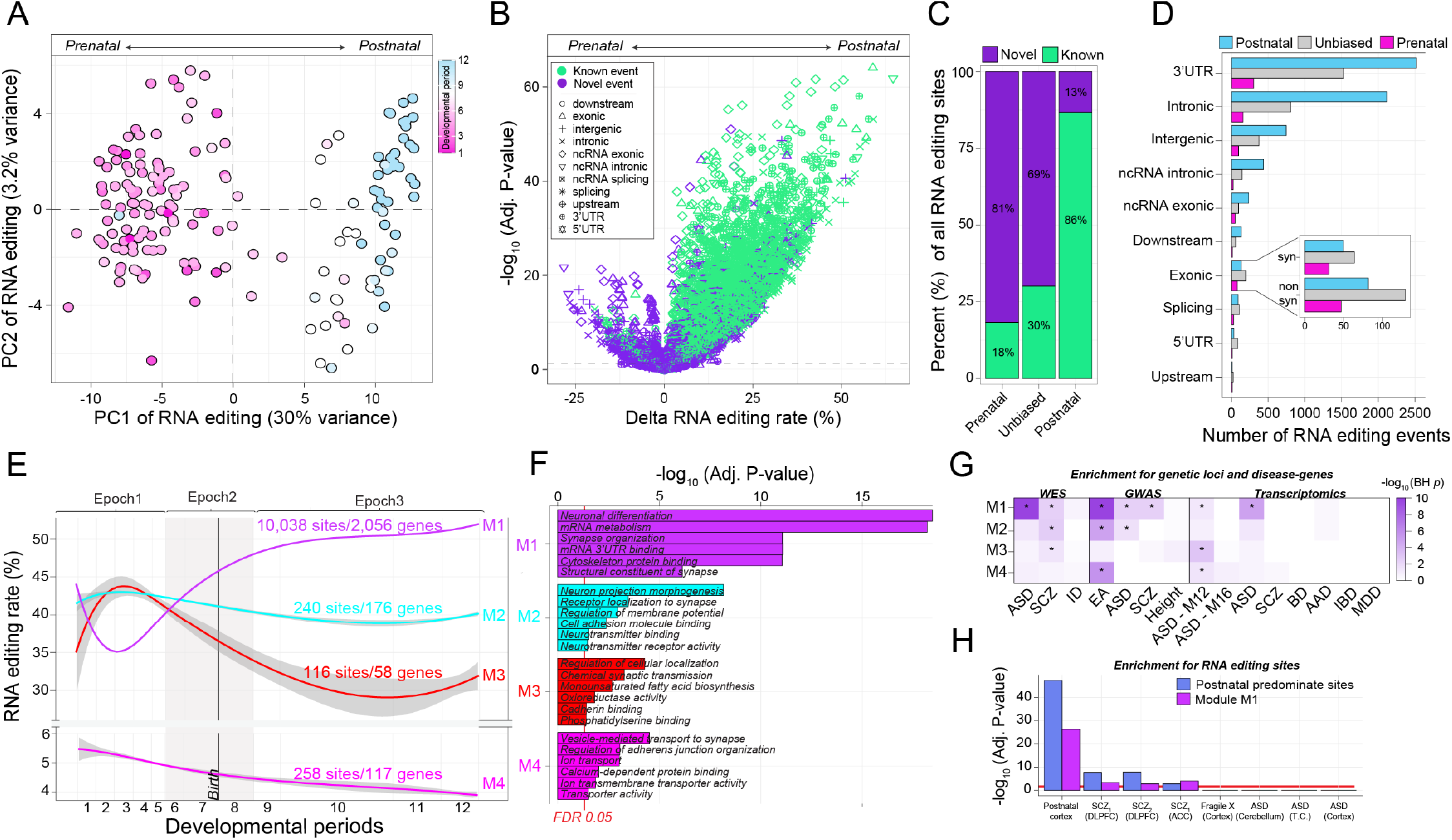
Developmental regulation of selective RNA editing sites. (**A**) Principal component analysis of RNA editing sites (N=10,652 sites) across all samples stratifies prenatal (pink) from postnatal (blue) periods. (**B**) Differential RNA editing analysis compares the strength of significance (-log_10_ FDR-adjusted *P*; y-axis) of developmentally regulated sites relative to delta editing rates (x-axis). Sites according to the developmental bias (prenatal, postnatal or unbiased) are partitioned according to (**C**) their novelty defined as presence/absence from existing RNA editing databases and (**D**) by genic region. (**E**) Weighted correlation network analysis partitioned all sites into four modules of sites with coordinated editing patterns across development. The average editing rates and standard error across all sites within each module (y-axis) are plotted across cortical development (log age, x-axis). (**F**) Each site was mapped to a gene and tested for functional annotation. The top 6 significant functional GO terms are displayed for each module. (**G**) Similarly, each module was examined for enrichment of neurodevelopmental disorder-related genes and gene sets identified from large-scale genetic and genomic studies. Color intensity reflects strength of significance (-log_10_ Adj. P). (**H**) Module M1 and all postnatal predominant sites were examined for enrichment of RNA editing sites found to be dysregulated in studies of postmortem brain tissue. Above the red line indicates a BH adjusted *p* < 0.05.

Most developmentally regulated editing sites increased in editing efficiencies and mapped to 3’UTRs (**Figure 4D**). Accordingly, we modeled whether RNA editing in 3’UTRs might influence miRNA binding efficiency (*see Materials and Methods*). A significant reduction in miRNA binding energy was observed in edited relative to un-edited 3’UTRs (*p*=2.3×10^−32^, Mann-Whitney U test) (**Figure S11**), suggesting a regulatory role for RNA editing in cortical development. Moreover, concordance analysis between RNA editing and corresponding gene expression profiles identified editing in 3’UTRs as having the largest overall effect and negatively correlated with gene expression across development (**Figure S12**).

To infer the biology of developmentally regulated events, unsupervised weighted correlation analysis identified four discrete groups of co-regulated editing modules (M) that varied in the number of editing sites, gene content and editing efficiencies (**Figure 4E, Figure S13**). All modules were enriched for general terms related to neuronal projection, synapses, axons and dendrites, but were also enriched for unique functional processes (**Figure 4F, Supplemental Table 3**). The majority of sites mapped to M1 (*n*^*sites*^= 10,038), which was postnatally biased (*R*=0.87, *p*=4.0×10^−56^), but implicated in biological processes known to be prenatally biased in gene expression, including neurogenesis, mRNA metabolism, cytoskeletal protein binding and 3’UTR binding. M2 (*n*^*sites*^= 204), the grey module, was prenatally biased (*R*=-0.78, *p*=2.0×10^−36^) and associated with neuronal projection, cell signaling, cell adhesion molecule binding and neurotransmitter binding and receptor activity. M3 (*n*^*sites*^= 116), the smallest module was also prenatally biased (*R*=-0.33, *p*=1.0×10^−5^) and implicated in synaptic transmission, fatty acid biosynthesis, oxidoreductase activity and cadherin binding. M4 (*n*^*sites*^= 258) was the module with the lowest level of editing activity, also prenatally biased (*R*=-0.73, *p*=2.0×10^−30^), and was implicated in processes related to ion transporter activity to synapse, regulation of adherens junction organization and calcium-dependent protein binding. Notably, M1 was significantly enriched for: *i)* genes that confer risk for autism spectrum disorder (ASD), schizophrenia and educational attainment; *ii)* differentially expressed genes in cortical samples from individuals with ASD; and *iii)* dysregulated RNA editing sites in cortical samples from individuals with schizophrenia and sites related to postnatal development (**Figure 4G-H**). Furthermore, supervised analysis of candidate gene sets confirmed that RNA editing had an overall negative association with neurogenesis, neuronal differentiation, mRNA metabolism (genes with high prenatal expression) and a positive association with synaptic signaling (genes with high postnatal expression) (**Figure S14**).

We validated the temporal trajectories for the vast majority of developmentally regulated sites in two smaller independent developmental brain transcriptome datasets, and observed high transcriptome-wide concordance of delta editing rates across forebrain (*R*^*2*^=0.58) and cerebellum development (*R*^*2*^=0.31) (**Figure S15**). Next, in 261 aged cortical samples, ∼90% of all common editing sites with sufficient coverage (*n*^*sites*^= 9,568) were reproduced and these data re-confirmed that M1 was consistently highly edited during advanced aging, whereas the remaining modules were edited to a lesser extent (∼0-2% editing efficiency) (**Figure S16A-C, Supplemental Table 4**). In addition, ∼20% of all common editing events (*n*^*sites*^= 2,168) were re-confirmed in hESCs and validated prenatally biased modules M2 and M3, which contained significantly more coverage and higher editing rates relative to modules M1 and M4 (**Figure S16D-F, Supplemental Table 4**).

### Dynamic developmental regulation of RNA recoding sites

RNA editing can recode amino acids and a number of recoding events play an important role for neuronal function and development. To expand upon this current knowledge, we observed that recoding sites displayed the smallest developmental changes in editing efficiencies compared to sites in other genic regions, as expected (**Figure 5A**). Second, recoding events were more highly conserved relative to sites in all other genic regions (**Figure 5B**) and enriched for components related to glutamate receptor complex (FDR *p*=4.3×10^−6^), ion-channel activity (FDR *p*=1.9×10^−3^), cadherin binding (FDR *p*=0.002), and mRNA metabolism (FDR *p*=0.002) (**Figure S17, Table S3**). Third, we identified 32 recoding events that displayed significant developmental changes in editing efficiencies (**Figure 5C-E**), including nine high-confidence novel recoding events that validate in advanced age (**Figure S18**). Ranking these sites by developmental effect sizes re-confirmed several recoding sites that are known to increase in editing efficiency through cortical development, including those mapping to a collection of excitatory, inhibitory and G-coupled protein receptors (*e*.*g*., *GRIK2, GABRA3, GRIA2, CYFIP2*). However, we also highlight numerous known recoding sites with previously undefined roles in cortical development: an arginine/glycine conversion (R/G) on the 3^rd^ exon of cyclin I (*CCNI*); an I/M and S/G site on the 3^rd^ exon of signal recognition particle 9 (*SRP9*); an R/G site on the 3^rd^ exon of SON DNA and RNA binding protein (*SON*, the cause of ZTTK syndrome); a K/R site on the 6^th^ exon of endonuclease VIII-like 1 (*NEIL*); a T/A site on the 6th exon of EEF1A lysine methyltransferase 2 (*EEF1AKMT2*), among others (**Figure 5D-E**). Finally, of the nine novel recoding events, seven were prenatally biased in their editing efficiencies and two were C-to-T edits, including a Q/R site on the 2^nd^ exon of ribosomal protein S6 kinase C1 (*RPS6KC1*), D/G site on the 11^th^ exon of MINDY lysine 48 deubiquitinase 3 (*MINDY3*), and an R/G site on the 12^th^ exon of sorting nexin 2 (*SNX2*), among others.

**Figure 5.**
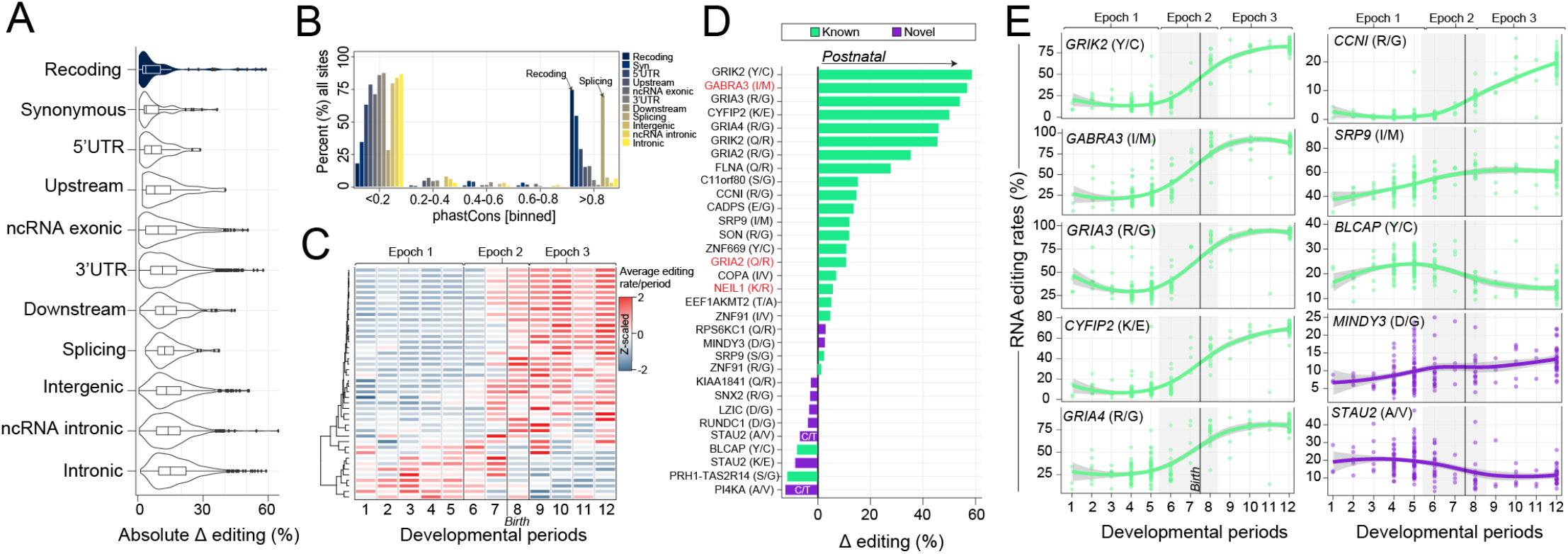
Temporal specificity of RNA recoding events. (**A**) Absolute delta editing rates parsed and ranked by genic region, where recoding events show the weakest developmental effect. (**B**) phastCons conservation metrics binned (x-axis) and the percent of all RNA editing sites per region are displayed per bin (y-axis). (**C**) Unsupervised clustering and heatmap depiction of 32 RNA editing recoding events that significantly change across development. (**D**) Ranking of 32 recoding sites by strength of developmental effect on editing rates (x-axis). Genes and sites in red indicate those where editing rates reach 100% and those in purple reflect novel recoding events. (**E**) Examples of editing rates (y-axis) for ten recoding sites across development, including five well-known sites (left), three known documented sites with unexplored roles in development (right) and two novel recoding (right) (x-axes). Loess curves were used to fit the data.

### RNA hyper-editing through human cortical development

Using BrainVar data, we first measured RNA hyper-editing across all substitution types and A-to-G accounted for ∼99% of edits (**Figure S19**). Therefore, this class was the primary focus of ensuing analyses. Hyper-editing in 3’UTRs and introns comprised ∼80% of all events with very few events in coding regions (∼0.24%). Hyper-editing occurred in clusters of ∼80bp in length and each cluster contains ∼10.8 editing sites (**Supplemental Table 5**). The rate of hyper-editing significantly increased from prenatal (µ=24,428 sites) through postnatal (µ=78,441 sites) periods (*p*=1.4×10^−17^) (**Figure 6A**). To minimize technical variability and enable a direct comparison of hyper-editing across development, we normalized the rate of hyper-editing to the number of million mapped bases per sample. Normalized hyper-editing rates increased in frequency into postnatal development (*p*=7.2×10^−17^, Cohen’s d=3.06) (**Figure 6B**). These rates were associated with the proportion of mature neurons (*R*^*2*^=0.41) and ADAR expression (**Figure S20**), and remained significantly postnatally biased following adjustment for neuronal cell type frequencies (*p*=1.0×10^−11^). The local sequence motif for all A-to-G hyper-editing sites was consistent with the expected distribution of guanosines upstream and downstream the target adenosine (**Figure 6C**).

**Figure 6.**
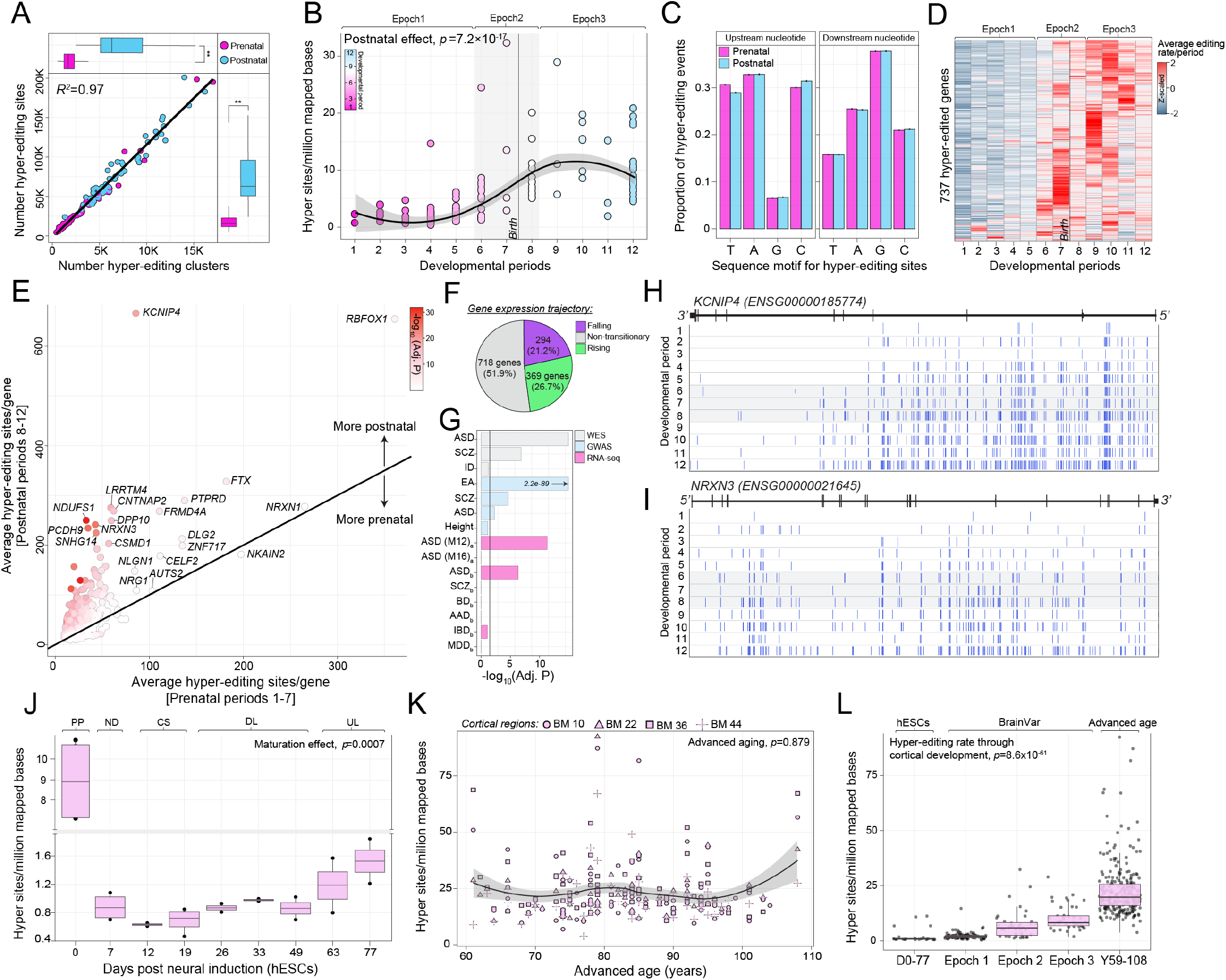
RNA hyper-editing signal through cortical development. (**A**) The number of hyper-editing sites (y-axis) is concordant with the number of hyper-editing clusters (x-axis). Significant increase (**, *p*<2×10^−16^) in the number of hyper-editing sites and clusters was observed between prenatal and postnatal periods (linear regression). (**B**) The number of RNA hyper-editing sites normalized by the number of uniquely mapped bases per sample. This normalized measure of hyper-editing (y-axis) was examined across 12 developmental periods (x-axis) prior-to and following adjustment for the proportions of mature neurons with linear regression (*p*=7.2×10^− 17^, *p*=1.0×10^−11^, respectively). (**C**) A local sequence motif for all hyper-editing events detected prenatally and postnatally are enriched for a depletion of guanosine -1bp upstream and an enrichment of guanosine +1bp downstream the target adenosine. (**D**) Heatmap depiction of genes that amass hyper-editing events into postnatal development. The number of hyper-editing events per period were averaged for each gene and z-scaled. (**E**) Average number of hyper-editing events per gene during prenatal periods 1-7 (x-axis) versus postnatal periods 8-12 (y-axis). Each gene is colored according to their FDR adjusted *p*-value significance. (**F**) The developmental expression trajectories for genes that amass A-to-G hyper-editing sites during postnatal periods. (**G**) Genes enriched for postnatal hyper-editing enrich for neurodevelopmental disorder associated genes and gene-sets curated from large-scale genomic studies (Fisher’s exact test). RNA hyper-editing barcode plots illustrate when and where hyper-editing events amass in two educational attainment genes: (**H**) *KCNIP4* (ENSG00000185774) and (**I**) *NRXN3* (ENSG00000021645). Normalized hyper-editing rates are examined prior-to and following adjusting for neuronal proportions across (**J**) a hESC model of corticogenesis (*p*=0.0007, *p*=0.03, respectively), (**K**) advanced aging (*p*=0.8, *p*=0.2, respectively) and (**L**) integrating hECS, BrainVar, and advanced aging (*p*= *p*=8.6×10^−61^, *p*=0.004, respectively). Note, cells during pluripotency (PP) were not included these analysis as they represent extreme hyper-editing outliers.

We identified 1,754 genes that amassed a significant number of A-to-G hyper-editing sites throughout development. After adjusting for the proportion of mature neurons and gene length, we retained 737 genes enriched with postnatal hyper-editing signal (**Figure 6D-E, Supplemental Table 5**). Approximately 74% displayed developmental expression profiles that are either falling or non-transient over development, suggesting that the hyper-editing enrichment is largely independent of the corresponding gene expression levels (**Figure 6F**). Genes enriched with postnatal hyper-editing signal were also enriched for genes implicated in genetic risk for autism spectrum disorder, educational attainment, and schizophrenia, as well as genes that are significantly differentially expressed in the cortex of individuals with ASD (**Figure 6G**). For example, gradual accumulation of hyper-editing events over cortical development was observed in the 1^st^ and 2^nd^ introns of potassium voltage-gated channel interacting protein 4 (*KCNIP4*) and in the 4^th^, 10^th^ and 13^th^ introns of neurexin 3 (*NRXN3*). (**Figure 6H-I**).

These findings were compared with hyper-editing signatures during *in vitro* corticogenesis and in cortical samples from advanced ages (**Supplementary Table 5**). In hESCs, a significant number of hyper-editing sites accumulated throughout neuronal maturation and differentiation (*p*=0.007) (**Figure 6J**). Notably, the frequency of hyper-editing was significantly elevated (µ=21,796 sites) during pluripotency, in contrast to the remaining days post neural induction (µ=2,163 sites, *p*=1.9×10^−9^), and a marked ∼58% of these sites mapped to coding regions (**Figure S21**). In cortical samples during advanced age, the normalized hyper-editing rate was not dynamically regulated (**Figure 6K**) but increased twofold compared to the Epoch 3 of BrainVar (*p*=5.7×10^−11^). All hyper-editing sites were enriched for a common sequence motif (**Figure S21-22**). Moreover, genes displaying a significant postnatal enrichment of RNA hyper-editing sites in BrainVar were validated during advanced age with high concordance (*R*^*2*^=0.58) (**Figure S23**). Taken together, we observe a substantial enrichment of A-to-G hyper editing events that accumulate in the cortex through development (**Figure 6L**).

Finally, given that the majority of hyper-editing sites accumulate in introns of neurodevelopmental genes, we tested whether these sites (*i*) are predicted to be splice altering, (*ii*) occur in retained introns and/or (*iii*) occur in ultra-conserved non-coding elements (UCNEs), which can act as transcriptional regulators. First, the number of intronic RNA hyper-editing events predicted to be splice altering increased throughout development (*p*=2.9×10^− 19^), but ultimately accounted for a small subset of the total number of hyper-editing sites (**Figure S24**). Second, while intron retention (IR) was more commonly detected in postnatal periods (∼57,311 introns, ∼10,848 genes) relative to prenatal periods (∼54,638 introns, ∼10,176 genes) (*p*=0.0006), the percent intron inclusion ratio was moderately elevated during prenatal periods (*p*=0.01) (**Figure S25A-D, Supplemental Table 6**). Importantly, ∼80% of genes enriched with postnatal intronic hyper-editing events display heightened IR during postnatal periods, but greater IR was not correlated with the expression of their host genes (**Figure S25E-F**). Third, a significant over-representation of postnatal hyper-editing signal was also detected in genes harboring UCNEs (*p*=6.3×10^−5^), which might underlie suboptimal regulation during ageing (**Figure S23C**).

### Conservation of increased selective editing and hyper-editing in mammalian cortical development

We tested whether these developmental patterns of RNA editing constitute a conserved regulatory mechanism involved in cortical physiology in two animal models of cortical development: *i*) bulk tissue RNA sequencing of four cortical regions of 26 rhesus macaques, from 60 post-conception days to 11 years; *ii*) whole-cortex RNA-sequencing of 18 wild-type mice, from embryonic day 14.5 to 21 postnatal months (**Supplemental Table 7**). In macaque, we observed a significant postnatal bias in both the AEI (*p*=3.5×10^−17^) and in the frequency of RNA hyper-editing sites (*p*=7.1×10^−10^) (**Figure 7A-C**). Hyper-editing clusters ranged in size from 65-72bp in length, each contained ∼7.8 hyper-editing sites, and all sites shared a local sequence motif similar to that observed in humans (**Figure 7D**). These findings also reproduced in mouse, including the increased rate of selective RNA editing (*p*=1.9×10^−12^), increased frequency of RNA hyper-editing sites (*p*=1.8×10^−7^), and conservation of common local sequence motif for all A-to-G hyper-editing events (**Figure 7E-H**). Furthermore, several developmentally regulated recoding sites validated across species, with few sites showing human-specific developmental regulation (**Figure S26**). Finally, we observed that global editing rates and the frequency of hyper-editing sites are highest in human cortex, followed by macaque and subsequently mouse (**Figure 7I**). Importantly, all hyper-editing sites in the current study were robust to potential false positives and were not confounded by common genomic variation (**Figure S27**).

**Figure 7.**
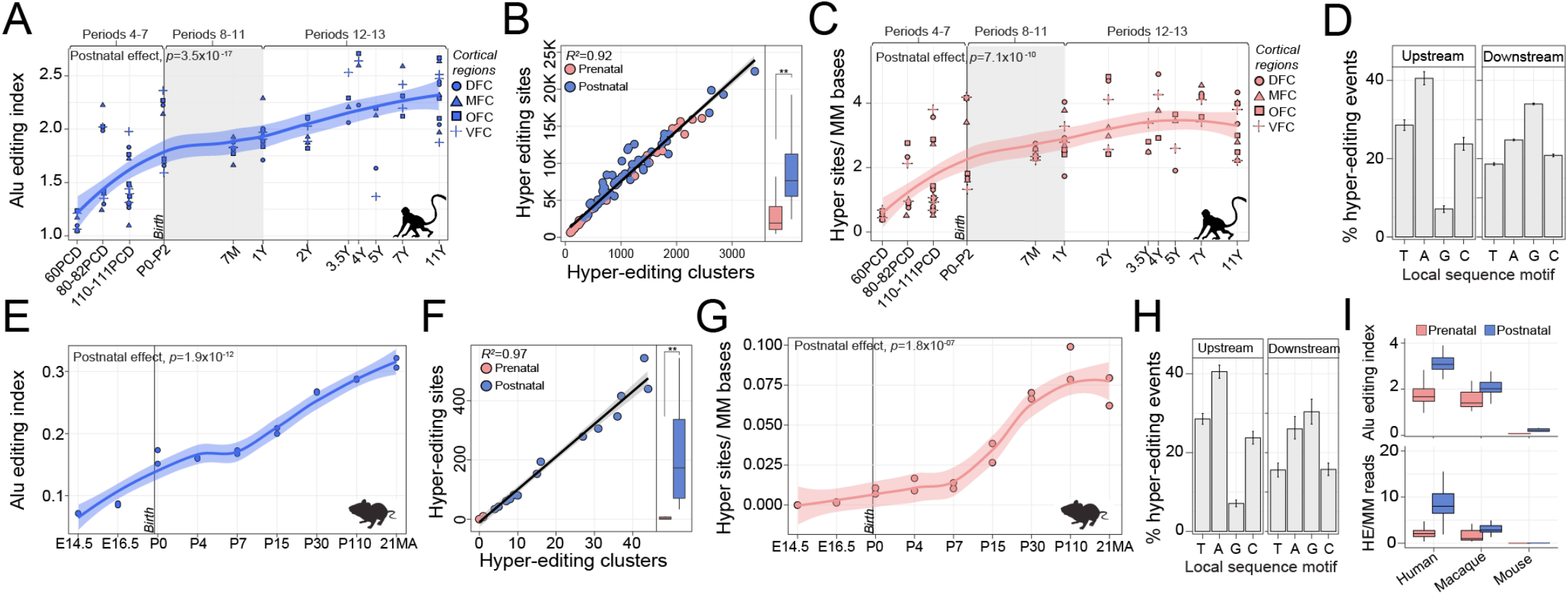
Evolutionary conservation of global editing in animal models of cortical development. (**A**) The AEI (y-axis) was computed for four cortical regions (DFC, MFC, OFC, VFC) in rhesus macaque across prenatal and postnatal periods (x-axis). Macaque developmental periods were matched with those closest to human (shown above) as previously shown (Y Zhu et al., 2018). (**B**) The number of hyper-editing sites (y-axis) is concordant with the number of hyper-editing clusters (x-axis) and the number of hyper-editing sites increases from prenatal to postnatal periods (**, indicates *p*=2.8×10^−6^). (**C**) Normalized hyper-editing rates across all developmental periods and cortical areas (*p*=7.1×10^−7^). (**D**) A local sequence motif for all hyper-editing events detected prenatally and postnatally are enriched for a depletion of guanosine -1bp upstream and an enrichment of guanosine +1bp downstream the target adenosine. (**E**) The AEI (y-axis) for whole cortex in mouse across nine developmental periods (x-axis). (**F**) The number of hyper-editing sites (y-axis) is concordant with the number of hyper-editing clusters (x-axis). Significant increase in the number of hyper-editing sites was observed between prenatal and postnatal periods (**, indicates *p*<2×10^−16^). (**G**) Normalized hyper-editing rates across all developmental periods. (**H**) A local sequence motif for all hyper-editing events detected are enriched for a depletion of guanosine -1bp upstream and an enrichment of guanosine +1bp downstream the target adenosine. (**I**) Comparison of the AEI and normalized hyper-editing rates across prenatal and postnatal stages between humans (using BrainVar), rhesus macaque and mouse. Linear regression was used to compute significance in all aforementioned tests.

### Developmental cis-edQTLs and co-localization with neurological traits and disorders

WGS data were used to detect SNPs that could have an effect on RNA editing levels (edQTL, editing quantitative trait loci). RNA editing levels were fit to SNP genotypes, covarying for developmental period, sex, the first five principal components of ancestry, as well as *ADAR1* and *ADAR2* expression. This permitted identification of editing levels under tight genetic control, irrespective of the corresponding ADAR activity. To distinguish temporal-predominant edQTLs, we performed three *cis*-edQTL analyses: i) prenatal samples only (*n*=116, periods 1-7); ii) postnatal samples only (*n*=60, periods 8-12); and iii) all samples (*n*=176, periods 1-12). We defined a 1Mb window (±) to search for SNP-editing pairs of an editing site and identified 10,803 cis-edQTLs from all samples, 5,386 cis-edQTLs from prenatal samples, and 1,008 cis-edQTLs from postnatal samples at FDR < 5% (**Supplemental Table 8**). These edQTLs comprised a total of 1039, 1,008 and 802 unique editing sites (eSites) across all samples, prenatal samples and postnatal samples, respectively, and each lead SNP was located close to their associated editing site (15kb±nt) (**Figure 8A-B**).

**Figure 8.**
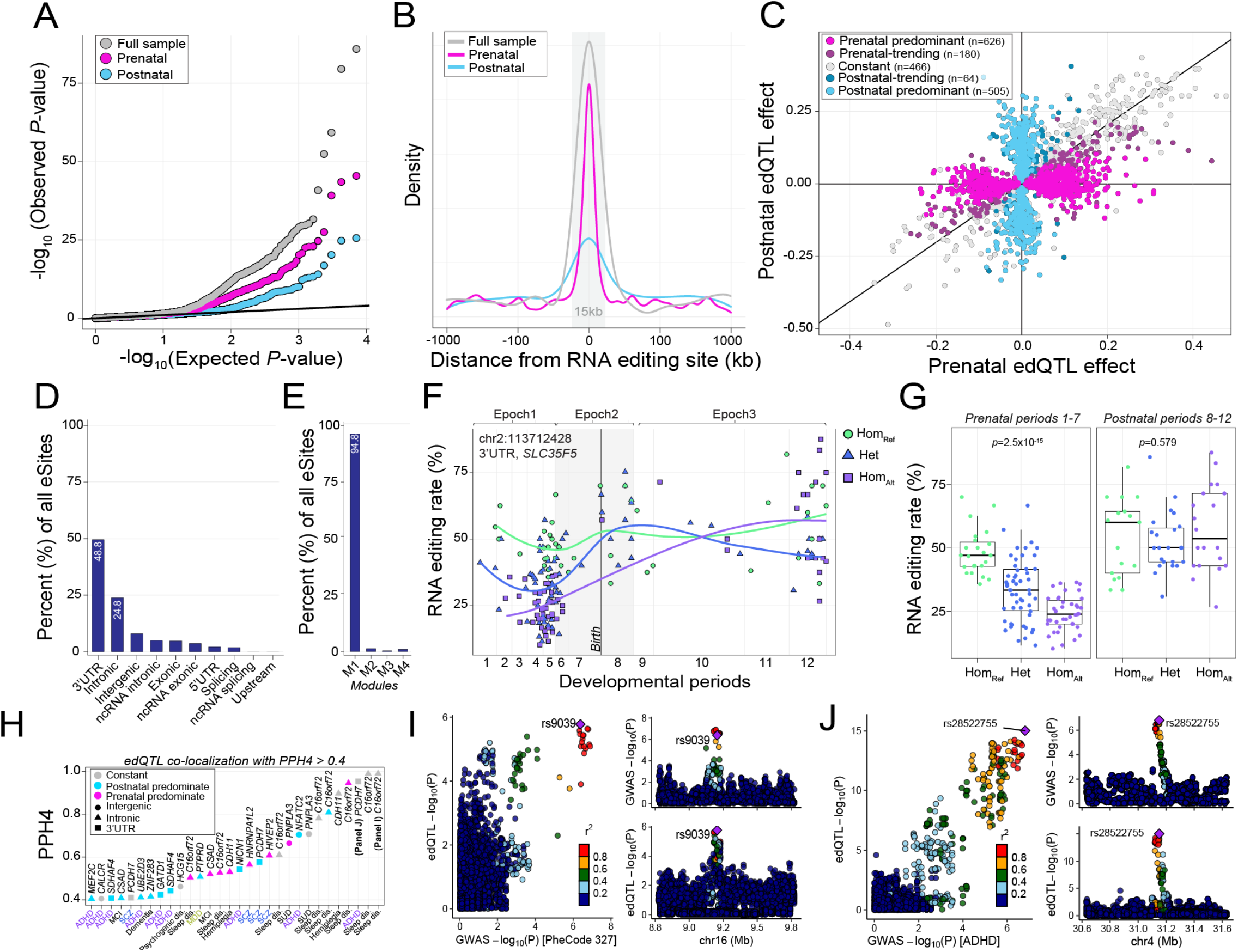
Cis-genetic regulation of RNA editing and co-localization with disease risk loci. (**A**) Quantile– quantile plot for association testing genome-wide *P* values between WGS-derived genotype dosages and 10,652 RNA editing sites across the full sample (*N=*176), prenatal periods 1-7 (N=116) and postnatal periods 8-12 (N=60) (linear regression and FDR correction via fastQTL). (**B**) Distribution of the association tests in relation to the distance between the editing site and variant for max-edQTLs. The grey box indicates ±15 kb relative to the editing site. (**C**) Prenatal (x-axis) and postnatal (y-axis) effects for the edQTLs with the smallest p value for all eSites (points). The edQTLs are split into five categories based on temporal predominance using effect size and statistical thresholds; categories are represented by color. The percentages (y-axis) out of all eSites according to (**D**) genic regions and (**E**) modules (M) of coordinated editing sites. (**F**) RNA editing rates binned by genotype for a top prenatal-predominant edQTL for *SLC35F5*. Lines represent loess trajectories for RNA editing in samples with each of three genotypes. (**G**) Boxplots for prenatal (left) and postnatal (right) periods with each of three genotypes. (**H**) Summary of co-localization results for edQTLs with a posterior probability of H4 (PPH4; y-axis) greater than 0.4. Colors reflect temporal predominance of the edQTL effect and shapes indicate the RNA editing site genic region across all UKBiobank neurological traits (in black) and large-scale GWAS (colored). Locuscompare plots of the top co-localized hit for (**I**) variant rs9039 associated with sleep disorders (PheCode 327) and intronic A-to-G editing site in *C16orf72* (chr16:9110931) and (**J**) variant rs28522755 associated with ADHD and 3’UTR A-to-G editing site in *PCHD7* (chr4:31143136).

The majority of edQTLs were consistent across development, reaching nominal significance in all three analyses with the same direction of effect (∼77%, **Figure 8C**). Many edQTLs were prenatal-predominant, with greater prenatal than postnatal effect sizes, while fewer edQTLs were postnatal-predominant, with significantly greater postnatal than prenatal effect sizes. Regardless of their temporal predominance, a substantial fraction of all corresponding eSites increased in editing efficiency through development (∼95%), enriched in module M1, and mapped to 3’UTRs and introns (**Figure 8D-E**). edQTL SNPs (edSNPs) were also examined for association with gene expression levels by calculating the overlap between edQTLs and previously computed expression QTL (eQTL) summary statistics. Overall, a total of 969 edSNPs (12%) were also associated with variation in gene expression, for which 27% were associated with a gene and one or more editing sites within the same gene.

Finally, we queried consistent and temporal predominant edQTLs for co-localization with common genetic risk variants for CNS disorders by leveraging genome-wide association studies (GWAS) summary statistics of four major neuropsychiatric disorders and several neurological traits from the UK BioBank. We found moderate co-localization (*PPH4* 0.3-0.8) for 98 loci across 15 traits and disorders and strong evidence of co-localization (*PPH4* > 0.8) for 9 loci across 4 traits and disorders (**Supplemental Table 9, Figure 8F**). The majority of disease variants co-localized with postnatal predominant edQTLs (*n*=72) followed by consistent (*n*=20) and prenatal predominant edQTLs (*n*=15). Postnatal edQTLs uniquely co-localized with variants that implicate risk for mild cognitive impairment and dementia while prenatal edQTLs uniquely co-localized with variants that implicate risk for generalized seizures and hemiplegia (**Supplemental Table 9**). Both consistent and prenatal predominant edQTLs co-localized with sleeping disorder traits, including two editing sites in the third intron of *C16orf72* (*PPH4*=0.99, *PPH4*=0.95, respectively) (**Figure 8H-I**). Notably, we also observed significant co-localization, with similar direction of effect, for a risk variant for attention deficit hyperactivity disorder (ADHD) with a prenatal predominant edQTL linked to an editing site in the 3’UTR of *PCDH7* (*PPH4*=0.95) (**Figure 8J**). The same edQTL also co-localized with a separate variant associated with schizophrenia (*PPH4*=0.57).

### Discussion

This work highlights RNA editing as an important, and relatively underexplored, species-conserved mechanism involved in cortical development. We provide a global temporal picture of RNA editing in the developing human cortex, from early progenitor cells to mature cells in centenarians. We recapitulated RNA editing profiles in non-human primate and murine models of neurodevelopment, underscoring RNA editing as a conserved evolutionary regulatory mechanism in mammalian cortical development. While these data do not immediately lend themselves to quick mechanistic interpretation, at each level of analysis we provide tangible examples and proofs-of-principle that establish starting lines for investigations to link RNA editing with mechanisms of cortical development.

Our analysis is anchored around BrainVar data, permitting precise base-specific quantifications of RNA editing from RNA-seq data guided by paired WGS (**Figure 1**). We observed that the late-fetal transition in gene expression between mid-fetal development and infancy is synchronized with the peak of RNA editing, that involves over 10,000 RNA editing sites that increase in editing efficiency throughout cortical development (**Figure 3**). This represents a substantial departure from the current perception of RNA editing in brain development^27-32^. Our results, akin to other recent reports^33,34^, indicate that these developmental periods are marked by clear changes in the proportions and compositions of various cell types in the brain, including the distinct transcriptional and RNA editing programs within these cell subsets. Recognizing the significance of this, we accounted for the proportion of mature neurons for each of our analyses, which preserved the significance of our findings and together indicated an increased rate of editing per cell through development. Providing these editing sites are involved in processes related to neuronal differentiation, mRNA metabolism and binding of actin cytoskeleton proteins and 3’UTR elements (**Figure 4**), alterations or reversals of these editing rates in the context of neurodevelopmental disorders, like schizophrenia (**Figure 4H**), may provide avenues for understanding their potential pathological impact.

Likewise, we uncover numerous temporally regulated recoding sites enriched for RNA metabolism and binding of cadherin and glutamate as well as genes abundant around the postsynaptic membrane (**Figure 5**). These results confirm several previously described recoding sites known to regulate Ca^2+^ permeability (*GRIK2*, Y/C, ∼58% increase)^16,17^, remodel actin cytoskeletal at excitatory synapses (*CYFIP2*, K/E, ∼50% increase)^19,20^ and guide gating kinetics of inhibitory receptors (*GABRA3*, I/M, ∼56% increase)^21,22^, but also highlight several sites with unexplored roles in neurodevelopment. For example, the Q/R site in *FLNA* increases ∼27% through development, a gene that anchors membrane receptors to the actin cytoskeleton and regulates their precise location and transport. The R/G site in cyclin-I (*CCNI*), a gene characterized by dramatic periodicity in protein abundance through the cell cycle and important for the immune response^37^, and the E/G site in calcium-dependent secretion activator (*CADPS*), previously shown to facilitate the rapid release of catecholamine and promote dense core vesicle exocytosis^38^ both increase in editing efficiency by ∼15%. However, the functional roles of these sites in cortical development have yet to be studied. We also identified seven novel recoding sites with high prenatal editing rates in regions of the genome with high mappability, which were not confounded by sequencing or other genome artifacts. These include a K/E site (chr8:73709034) in the double-stranded RNA binding motif of staufen double-stranded RNA binding protein 2 (*STAU2*). This gene is widely expressed in the brain, with high early fetal expression and is required for microtubule-dependent transport of neuronal RNA from the cell body to the dendrite^39^. The D/G site (chr1:9930448) in the beta-catenin-interacting protein domain of leucine zipper and *CTNNBIP1* domain containing gene (*LZIC*), may play a role in early developmental binding and regulation of beta-catenin^40,41^. A Q/R site (chr1:213071037) in the PX domain Ribosomal Protein S6 Kinase C1 (*RPS6KC1*), a phosphoinositide-binding structural domain involved in targeting of proteins to cell membranes^42^. Collectively, the recoding sites defined here have yet to be studied thoroughly in the context of neuronal maturation and cortical development and highlight numerous entry points for functional investigation.

RNA hyper-editing signal dramatically increased in frequency through development into advanced aging. Stretches of hyper-editing were predominantly found in Alu rich regions of the genome and more specifically, within introns and 3’UTRs that putatively form double-stranded RNA structures, consistent with recent reports^43-45^. Our analysis refined the developmental timing of these events and identified hundreds of specific genes that accumulate hyper-editing sites through development, most of which occur within retained introns of ion-channels and genes essential for typical development (**Figure 6**). Most hyper-edited regions are seldom detected as non-edited and thus may represent key ADAR targets. Markedly, hyper-editing events amassed in ultra-conserved non-coding segments, sequences that exhibit extremely high conservation across vertebrate genomes and cluster in non-coding regions of genes coding for transcription factors and key developmental regulators^46,47^. Thus, build-up of age-related RNA hyper-editing in these regions, and corresponding gene products, might underlie their suboptimal regulation and activity during aging, contributing to senescent age-related phenotypes.

While ADAR enzymes catalyze A-to-I RNA editing, the degree to which common genetic variation sets the mode and tempo of editing levels through development is a novel line of enquiry. We identified hundreds of edQTLs that have a greater effect on editing levels prenatally or postnatally (**Figure 8, Supplemental Table 8-9**). When combined with findings from large-scale GWAS, consistent and temporal-predominant edQTLs co-localized with a total of several neurological traits and disorders, including sleep disorders, ADHD, schizophrenia and cognitive impairment. Given that most risk loci reside in non-coding regions, the vast majority of edQTLs and genetic colocalization occurred with RNA editing sites in 3’UTRs, and did occur in coding regions or with RNA recoding events. We identified ADHD risk variant rs28522755 (chr4: 31143818 A/G)^48^ co-localized with a consistent edQTL linked to an A-to-G editing site in the 3’UTR of *PCDH7*. A separate postnatal predominant edQTL, linked to the same 3’UTR editing site in *PCDH7*, also co-localized with risk variant rs13145415 (chr4:30823157 T/G) for schizophrenia^49^. *PCDH7* plays a significant role in axonal guidance, predominantly through neuronal cell adhesion and short range signaling, and it has been proposed that mutations in *PCDH7* could introduce alterations in neuronal localization and perhaps brain structure^50^. Taken together, these two variants might exert their adverse consequences through altered 3’UTR RNA editing, shaping *PCDH7* gene expression and function, potentially offering new therapeutic avenues. Notwithstanding sample size limitations, these results identify temporal predominant edQTLs and GWAS-edQTL co-localization, thereby generating specific hypotheses for the role of RNA editing regulation in cortical development and neurological disease etiology.

In sum, RNA editing acts as conserved epitranscriptional mechanism in cortical physiology and is characterized by a global expansion of RNA sequence diversity. RNA editing rates increase throughout neurodevelopment, are commonly genetically regulated, and occur disproportionally in 3’UTRs of neurodevelopmental genes. This work further highlights variation in RNA editing as an intermediate mechanism that influences phenotypes of complex neurological traits and disorders. Collectively, these results serve as a resource toward classifying functionally relevant editing sites in the brain, as they uncover highly developmentally and genetically regulated sites which can serve to bridge the gap between their function and regulation of complex traits.

## Author contributions

Conceptualization: MSB

Data analysis: MSB, RC, EM, LS, LL, XF, MW, JW

Visualization: MSB, RC

Funding acquisition: MSB, SS, BZ, BD, KR, NS, JB

Supervision: MSB, BZ, BD, KR, JB, NS, SS

Writing – original draft: MSB, RC, LS, LL, XF, MW, BZ, JW, BD, KR, JB, SS

## Data and materials availability

The majority of data are available in supplementary tables. All code and larger data tables and files are available at GitHub (https://github.com/BreenMS).

