## Supplemental Methods and Figures for "Expansion of RNA sequence diversity and RNA editing rates throughout human cortical development"

[‡]*Individuals share second authorship.*

### Materials and Methods

***Tissue samples and data types used in the current study***

Multiple RNA-seq datasets were harnessed to quantify RNA editing across a range of developmentally distinct samples. All data were run through the same RNA-seq and RNA editing processing pipeline to obtain comparable results (see Processing pipeline). Raw FASTQ files from each of the following datasets were downloaded from either the sequencing read archive or Synapse.org:

1. **Human cortical development**: BrainVar (syn21557948) included 176 paired-end (100bp) samples of human dorsolateral prefrontal cortex (DLPFC) covering 12 distinct developmental periods. Processed whole-genome sequencing data were also leveraged for these matching individuals. The frontal cerebral wall was assayed in nine brains prior-to ten post-conception weeks.
2. ***In vitro* corticogenesis (hESCs)**: CORTECON (GSE56796) included 24 single-end (50 bp) samples of human cerebral cortex development from hESCs across nine time-points (days 0, 7, 12, 19, 26, 33, 49, 63, and 77).
3. ***In vitro* corticogenesis (hiPSCs)**: A total of 127 paired-end (100 bp) samples from a human induced pluripotent stem cell (hiPSC) neuronal differentiation time course (PRJNA596331) covering early differentiating cells (days 2-9), neural progenitor cells (day 15), assembled neuroepithelial rosettes (day 21) and more differentiated neurons (days 49-77).
4. **Advanced aging**: Mount Sinai Brain Bank (MSBB; syn7416949) included 261 single-end (100 bp) samples covering four cortical regions, including BM10, BM22, BM36 and BM44. This subset of samples was analyzed as they were largely free of plaque and tangle neurocognitive pathologies.
5. **Rhesus macaque** (*Macaca mulatta*): A non-human primate model of neurodevelopment (PRJNA448973) was leveraged, covering 96 single-end (75bp) samples, comprising 26 unique donors and four prefrontal cortical areas: MFC – medial prefrontal cortex; OFC – orbital prefrontal cortex; DFC – dorsolateral prefrontal cortex; VFC – ventrolateral prefrontal cortex.
6. **Mouse** (*Mus musculus*): A murine model of neurodevelopment (SRP055008) was leveraged including 18 paired-end (200 bp) cortical samples at nine time points: embryonic day 14.5 (E14.5), E16.5, postnatal day 4 (P4), P7, P17, P30, 4 months, and 21 months.
7. **Single-cell RNA-sequencing**: Two studies were leveraged to query ADAR expression in single cells. The first sampled the fetal developing cortex from 6 to 37 post-conception weeks (phs000989.v3,

https://cells.ucsc.edu/?ds=cortex-dev) and the second sampled three adult postnatal DLPFC tissues (syn15672826).

8. **Human organ development:** A total of 315 single-end (100 bp) samples from seven human tissues sampled across prenatal and postnatal periods (4 PCWs-59+ years) were downloaded from ArrayExpress (E-MTAB-6814). Four samples proved problematic and a total of 311 samples were included in subsequent analyses: bulk brain (*n*=55), heart (*n*=48), cerebellum (*n*=55), kidney (*n*=40), liver (*n*=50), testis (*n*=41) and ovary (*n*=18). Notably, ovary was sampled during fetal periods only.

#### *RNA editing site detection and annotation*

A graphical overview of our RNA editing pipeline is displayed (right). All FASTQ files were mapped to human reference genome using STAR[51] v2.7.3 and the following parameters were optimized: chimSegmentMin=15; chimJunctionOverhangMin=15; outSAMstrandField=intronMotif. For each sample, this produced a coordinate-sorted BAM file of mapped reads, including those spanning splice junctions. RNA editing sites were quantified *de novo* from sorted bam files using reditools v2.0[52] and the following parameters: -S -s 2 -ss 5 -mrl 50 -q 10 -bq 20 -C -T 2 –os 5. All analyses considered read strandedness (-s) when appropriate. A number of filtering steps were applied to retain only high-quality, high-confident bona fide RNA editing sites: i) all multi-allelic events were split into independent classes; ii) a minimum coverage of 10 reads; iii) a minimum coverage of 3 edited reads; iv) any sites mapping to homopolymeric regions were discarded; v) any sites mapping to high confidence heterozygous or homozygous genomic calls (as in the case with BrainVar data) were discarded; vi) any sites mapping to common genomic variation in dbSNP(v150) and those in gnomAD with minor allele frequency greater than 0.05 were discarded; vii) remaining sites were annotated using ANNOVAR[53] to gene symbols using RefGene, repeat regions using RepeatMasker v4.1.1, known RNA editing sites using the most recent version of REDIportal[54] and conservation metrics were gathered from the *phastConsElements30way* table of the UCSC Genome Browser, which consists of multiple alignments of 30 vertebrate species and measurements of evolutionary conservation using *phastCons* and *phyloP* from the PHAST package[55]. Sites mapping to ultra-high signal artifact regions of the genome were flagged[56]. Subsequently, annotated sites were further filtered using the sample size of the cohort under investigation by requiring sites to have: vii) a detection rate in at least 70% of the current sample, ix) a minimum editing rate of 5% or greater across all samples and x) no more than 20% missing values per sample. The resulting RNA editing data frames (whereby rows are editing sites and columns are samples) in this study contained no more than ~6% missing data on average. Any missing values were imputed using predictive mean matching method in the *mice* R package[57], using five multiple imputations and 30 iterations. All resulting sites from these steps were subsequently referred to as high-confidence selective RNA editing sites and were used for downstream analysis.

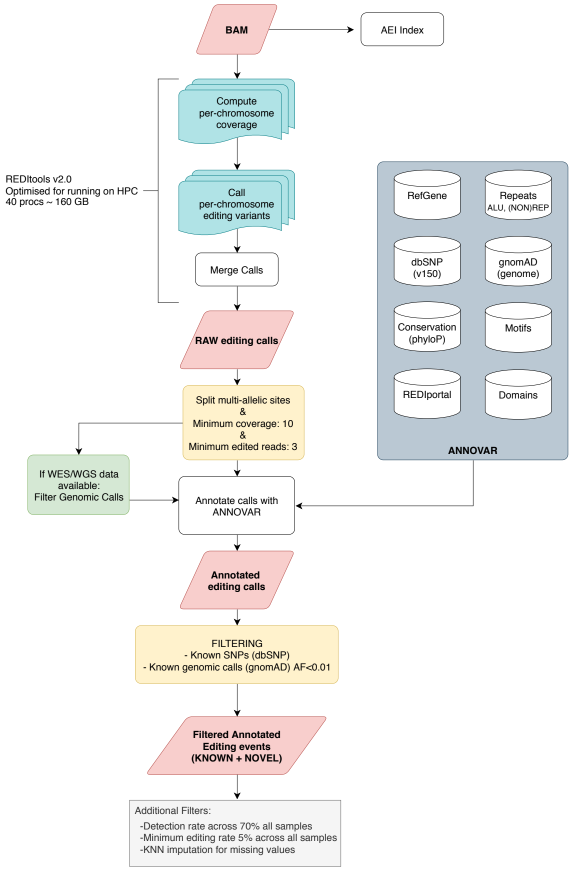

#### *Querying BrainVar RNA editing sites in independent data*

RNA editing sites in BrainVar were queried in the following independent datasets: hESC and hiPSC models of corticogenesis, human forebrain and hindbrain tissue across development, and in all MSBB samples during advanced aging. Nucleotide coordinates for all well documented editing sites in BrainVar were used to extract reads from each independent sample using the samtools mpileup function, as previously described[58]. This approach quantifies the total number of edited reads and the total number of un-edited reads that map to each RNA editing site detected in BrainVar, thereby producing a rich source of editing information across independent data.

#### *Computing an Alu editing index (AEI)*

The AEI method v1.0[59] was leveraged to computed the Alu editing index (AEI) using a STAR mapped bam file (described above) as input. The AEI is computed as the ratio of edited reads (A-to-G mismatches) over the total coverage of adenosines and is a robust measure that retains the full Alu editing signal, including editing events residing in low-coverage regions with a low false discovery rate. For human and macaque RNA-seq samples, we set our predetermined genomic regions to all SINE/Alu repeats using the Alu bed table of the UCSC genome browser defined by RepeatMasker. For mouse, we set these genomic regions to all B1-SINE and B2-SINE elements, where most of the editing activity in mouse occurs. Common genetic variation was also discarded for all species. Because we compare the AEI across independent studies and samples, it is important to emphasize that AEI has been shown to be highly scalable and comparable across postmortem brain RNA-seq samples generated across seven unique laboratory sites and five different library preparation protocols[59].

#### *Cellular deconvolution of bulk DLPFC tissue*

To identify changes in cellular composition in bulk DLPFC tissue, we applied non-negative least squares (NNLS) from the MIND R package[60] and utilized the Darmanis et al., [61] signature matrix which contained a mixture of six major cell types: mature (adult) neurons, immature (fetal) neurons, astrocytes, oligodendrocytes, oligodendrocyte precursor cells (OPCs) and microglia. NNLS, executed through the est_frac function, was applied to $\log_2$ count per-million (CPM) transformed data using the *limma* package in R[62]. We focus our predictions on these major cell types in an effort to reduce noise and to evaluate a distribution of cell type changes that reflect an approximate expected distribution based on prior work.

#### *Differential RNA editing analysis*

To identify sites with differing levels of RNA editing across cortical development, we implemented linear model though the *limma* R package[62] covarying for the possible influence of sex and ancestry. Notably, RNA integrity (RIN), postmortem interval (PMI) and pH explained < ~0.1% in RNA editing variance and were not included in the model. However, given that ADAR enzymes increase in expression through development and RNA editing is a process that is highly abundant in neurons, which also increase in frequency through development, we fit additional models to adjusted for both *ADAR1* and *ADAR2* expression (the enzymatically active editing enzymes) and the estimated neuronal composition for each individual. These adjustments enabled us to identify potential ADAR-dependent editing activity and to disentangle RNA editing activity changes from alterations in neuronal proportions. All significance values were adjusted for multiple testing using the Benjamini and Hochberg (BH) method to control the false discovery rate (FDR). Sites passing a multiple test corrected $P$-value $< 0.05$ were labeled significant.

#### *Co-editing network analysis and functional annotation*

We democratized editing rates across development by applying unsupervised weighted correlation network analysis[63]. One signed network was constructed for all BrainVar samples. To construct a network, the absolute values of Pearson correlation coefficients were calculated for all the possible editing site pairs and resulting values were transformed using a $\beta$-power of 7. The WGCNA cut-tree hybrid algorithm was used to detect RNA sites with coordinated editing activity, co-editing modules, with the following optimizations: minimum module size of 10 sites, tree-cut height of 0.9999 and a deep-split option of 4. Co-editing modules were functionally annotated for GO ontology terms relevant to cellular components, molecular factors, biological processes and metabolic pathways using the ToppFunn module of ToppGene Suite software[64]. We set a genomic background defined as all genes harboring at least one editing site and tested for significance using a one-tailed hyper-geometric distribution with a Bonferroni correction. This is a proportion test that assumes a binomial distribution and independence for probability of any gene belonging to any set. We use a one-sided test because we are explicitly testing for over-representation of genes that harbor editing sites across hundreds of GO categories, without any *a priori* selection of candidate gene sets. Notably, this enrichment analysis resulted in extensive lists of over-represented GO categories. Therefore, we applied REVIGO[65] using default parameters to summarize and remove redundant and similar terms from extended lists of GO terms using semantic clustering. These procedures for functional annotation were also carried out on all RNA recoding sites.

#### *Enrichment for disorder-related genes and RNA editing sites*

Co-editing modules were interrogated for over-representation of neurodevelopmental and neuropsychiatric disorder-related genes. Four tiers of gene sets were collected to examine overlap with co-editing modules: (1) whole exome sequencing (WES)-derived gene sets implicated in risk for autism spectrum disorder (ASD)[66], schizophrenia[67] and intellectual disability (ID)[68]; (2) genome-wide association study gene sets (gene(s) nearest candidate risk variants) that implicate risk for ASD[69], EA[70], schizophrenia[71] and height[72] (as a negative control); (3) transcriptome-derived gene sets of genes dysregulated in postmortem brain tissue of ASD[73], schizophrenia, bipolar disorder, major depressive disorder and alcoholism and inflammatory bowel disease[74,75] (as a negative control); (4) transcriptome-derived gene sets of RNA editing sites associated with postnatal cortical development[32], schizophrenia[58], Fragile X Syndrome and ASD[76]. To compute significance of all intersections, we used the GeneOverlap function in R[77] which uses a Fisher's exact test (FET) and an estimated odds-ratio for all pair-wise tests based on a background set of genes detected in the current study. When testing overlap across co-editing modules, all pairwise tests were adjusted for multiple testing using BH procedure to control the FDR.

#### *Identification and annotation of RNA hyper-editing sites*

RNA hyper-editing is defined as consecutive editing of many neighboring adenosines within an extended region or cluster. RNA reads that undergo extensive hyper-editing will not align to the reference genome due to the high load of mismatches. Thus, to identify hyper-edited reads in the current study, all discarded and unmapped reads following STAR alignment (described above) were converted to fastq format and used as input for hyper-editing analysis. We adopted a recently described RNA hyper-editing pipeline[43,44] with additional modifications, in brief:

(1) all unmapped reads with filtered by phred (<25), presence of simple repeat structures, high *N* content and fraction of a nucleotide per read being too great (>60%) or too low (<10%); (2) for resulting reads, all adenosines were transformed into guanosines in both the RNA sequences and genome reference sequence; (3) RNA reads were re-aligned using BWA-aln v0.7.15, all unmapped reads were discarded and subsequently all true adenosines were recovered to identify confident A-to-G edits; (4) processing and filtering of multi-mapped reads by selecting only the location with the largest fraction of A-to-G to all mismatches (providing this fraction was ≥10% higher than in all other locations, otherwise the read was discarded) and filtering of hyper-editing clusters to obtain high-quality (Phred ≥30) A-to-G mismatches in which the number of A-to-G mismatches was ≥5% of the read length and >60% (80% for read lengths ≤60 bp) of the total number of mismatches. This filtering also removed clusters that were too dense (>90% of read length), A-to-G mismatching contained within the first or last 20% of reads ends and clusters >60% of an individual nucleotide. Additional processing steps were applied to the resulting hyper-editing sites and clusters: (1) we extended cluster boundaries by the average distance between editing sites per cluster and subsequently merged clusters with overlapping coordinates (cluster length is a commonly product of read length); (2) all resulting hyper-editing sites were annotated using ANNOVAR (described above); (3) sites mapping to common genomic variation in dbSNP(v150) and those in gnomAD (maf>0.05) were discarded; (4) sites mapping to paired high-confidence private genomic calls were discarded. This entire process was reiterated to search for additional hyper-editing substitution types (*e.g.* A-to-C, G-to-C etc…).

To minimize the effect of sequencing batches and differences in library size, we normalized the total number of hyper-edited reads over the total number of uniquely mapped bases from the initial STAR alignment for each sample. To obtain the number of uniquely mapped bases, we ran Picard v2.22.3 (http://broadinstitute.github.io/picard/) on each mapped bam file. This enabled us to accurately compare normalized hyper-editing rates across independent studies, different developmental periods and different cortical regions.

***Differential analysis of gene-centric RNA hyper-editing***

To test for differences in the frequency of RNA hyper-editing per gene across development, we first queried genes that at least one RNA hyper-editing site in at least 70 BrainVar samples (~40%). This unsupervised measure was used to remove genes with too few RNA hyper-editing sites across all samples. All remaining missing values were treated as zero. Subsequently, we used the linear modelling framework of *limma*[62] to test for differences in the frequency of RNA hyper-editing events, while covarying for the possible influence of sex and ancestry. Additional models were fit that covaried for the proportions of mature neurons, *ADAR1* and *ADAR2* expression and gene length. To adjust for gene length, we repeated these analyses on a residualized matrix whereby the total number of edits per gene were normalized by the log gene length (geneEdits~log(gene length)). All significance values were adjusted for multiple testing using the Benjamini and Hochberg (BH) method to control the false discovery rate (FDR). Sites passing a multiple test corrected *P*-value < 0.05 were labeled significant.

***RNA editing and interactions with miRNA binding, alternative splicing and intron retention***

To compute differences of miRNA minimum free energy on edited and un-edited 3'UTRs, we used the following approach: (1) For each 3'UTR editing site, we obtained two 101bp sequences (50bp +/- the target editing site), where the only difference between these sequences was either an unedited (A) or edited (G) site; (2) We used miRANDA[78] to compute local alignments of 3'UTR sequences against all mature miRNAs obtained from miRBase; (3) miRANDA was used to compute the stability of the resulting RNA duplex and minimum free energy (DG kal/mol); (4) This process was re-iterated for high confidence alignments that occur in either in miRNA seed regions (setting --strict parameters) or any miRNA region. A Mann-Whitney U test was used to test for significance for differences in minimum free energy.

To compute the probability of intronic sites to be splice altering, we applied SpliceAI, a deep neural network that accurately predicts splice junctions from an arbitrary pre-mRNA transcript sequence[79]. We quantified all high-confidence intronic site-selective sites defined as prenatal, postnatal and commonly edited across development (Table S2), as well as all *bona fide* intronic RNA hyper-editing sites per donor across development. This approach enables precise prediction of non-coding RNA editing events that cause cryptic splicing and generates two outputs for each site: (1) Delta score, which is the probability of the RNA editing site being splice-altering and ranges from 0 to 1. In the primary publication of this work, a detailed characterization is provided for 0.2 (high recall), 0.5 (recommended), and 0.8 (high precision) cutoffs; (2) Delta position, which conveys information about the location where splicing changes relative to the variant position.

To compute the percent of intron retention over development, we applied Systematic Investigation of Retained Introns (SIRI)[80]. We provided mapped STAR bam files (as described above) as input together with strandedness (forward) and read length (100bp). To compute the percent of intron inclusion, inclusion was measuring using the PI junction field, defined by inclusion counts divided by the sum of inclusion and skipping junction counts. We selected only introns with a unique intron annotation (U introns) that are not involved in other alternative processing events. Introns subjected to PI measurement were also required have an intron length greater than and equal to 60 and have a sum of EE + EI + IE reads be greater than and equal to 20, as previously described[80].

#### *Identification of RNA editing quantitative trait loci*

Cis-edQTLs were identified for all high quality, common variants within 1 Mb (±) of an editing site using the fastQTL permutation based analysis[81] using a total of 1,000 permutations, with developmental period, sex, *ADAR1*, *ADAR2*, and the first five principal components of common variant ancestry as covariates. Associations between genotype dosages and period are measured with linear regressions. This analysis was run on three partitions of the BrainVar data set: the full sample (*n*=176, periods 1-12), prenatal samples only (*n*=112, periods 1-6), and postnatal samples only (*n*=60, periods 8-12). Separately for the results of each analysis, false discovery rate (FDR) was calculated for all gene-variant pairs using the Benjamini-Hochberg procedure. We then classified all edQTLs with FDR ≤ 0.05 from at least one analysis into one of five categories defined by the temporal specificity of their edQTL effects (as previously described[33]): (1) Constant edQTLs (consistent effects across development) are defined as edQTLs with FDR ≤ 0.05 in the complete sample analysis, same direction of effect and unadjusted p ≤ 0.05 in both the prenatal and postnatal analyses; (2) Prenatal-predominant edQTLs (strongest effects during prenatal development) are defined as FDR ≤ 0.05 in the prenatal analysis, unadjusted p > 0.05 in the postnatal analysis; (3) Postnatal-predominant edQTLs (strongest effects during postnatal development) are defined as FDR ≤ 0.05 in the postnatal analysis, unadjusted p > 0.05 in the prenatal analysis; (4) Prenatal-trending edQTLs were defined as those that did not fit into earlier categories, but had higher prenatal effects ($B_{Pre} > B_{Post}$); and (5) Postnatal-trending edQTLs were defined as those that did not fit into earlier categories, but had higher postnatal effects ($B_{Post} > B_{Pre}$).

#### *Co-localization with UK biobank GWAS summary statistics*

We leveraged GWAS summary statics for 109 neurological and mental health-related traits and disorders from the UKBioBank[82] (ftp://share.sph.umich.edu/UKBB_SAIGE_HRC/) along with GWAS summary statics for attention deficit hyperactivity disorder (ADHD)[48], schizophrenia[49], bipolar disorder[83], major depressive disorder[84]. For each set of summary statistics, genome-wide significant ($P<5.0\times10^{-8}$) loci were defined by linkage disequilibrium r2 > 0.6 start and end positions and edQTL sites overlapping those loci were considered for analysis. GWAS and edQTL summary statistics (beta, standard error) for SNPs within each GWAS locus were used as input to coloc2[85], and posterior probabilities for five hypotheses (H0, no GWAS or edQTL signal; H1, GWAS signal only; H2, edQTL signal only; H3, GWAS and edQTL signal but not co-localized; H4, co-localized GWAS and edQTL signals) were estimated for each locus. Loci with posterior probability for hypothesis H4 (PPH4) between 0.3-0.8 were considered to have moderate co-localization while PPH4 ≥ 0.8 was considered to demonstrate strong Bayesian evidence for co-localization.

*Supplemental Figures*

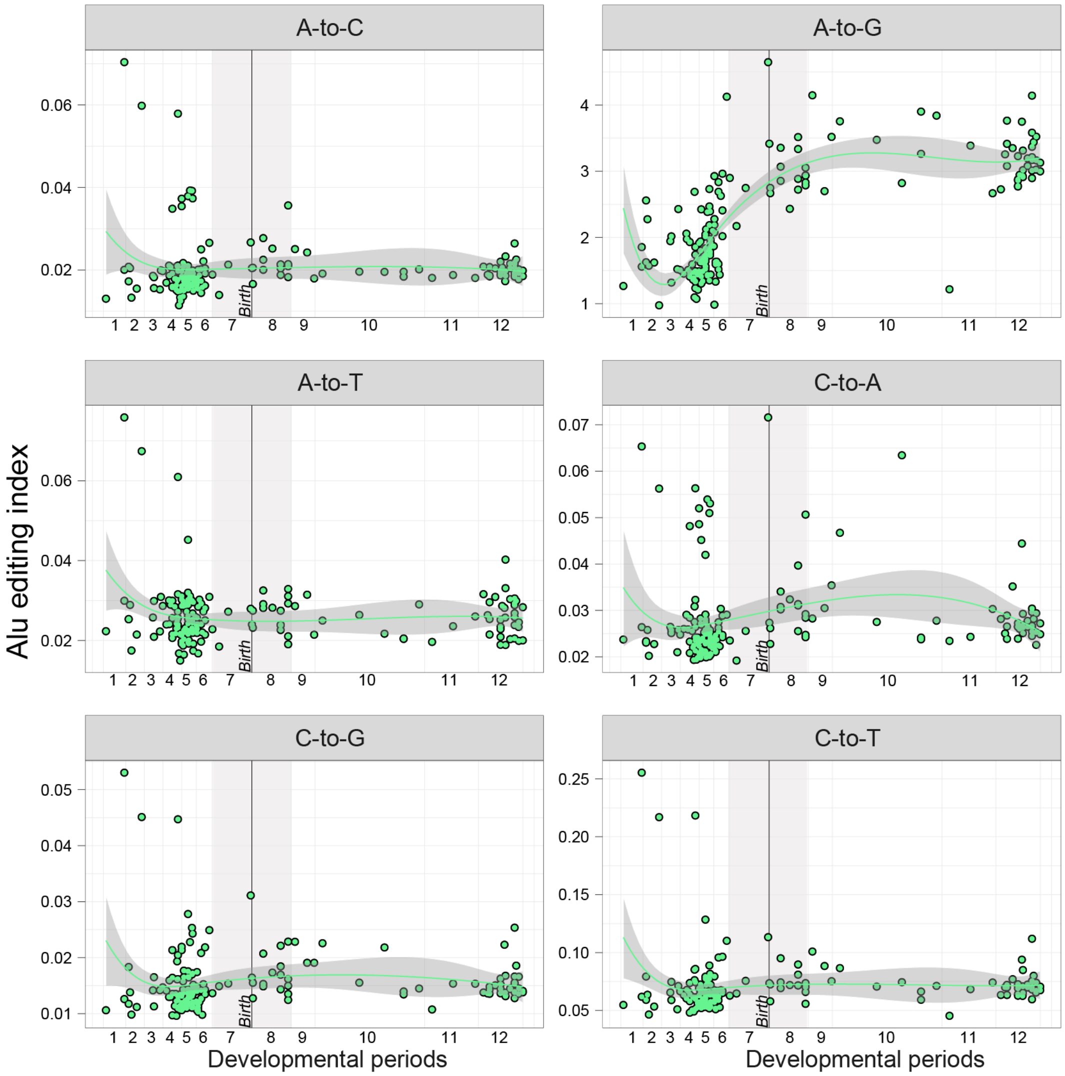


**Supplemental Figure 1. Alu editing index (AEI) by substitution types.** The AEI measured six different substitution classes for each BrainVar DLPFC sample (A-to-C, A-to-G, A-to-T, C-to-A, C-to-G, C-to-T; y-axes) across all 12 developmental periods (log age, x-axes).

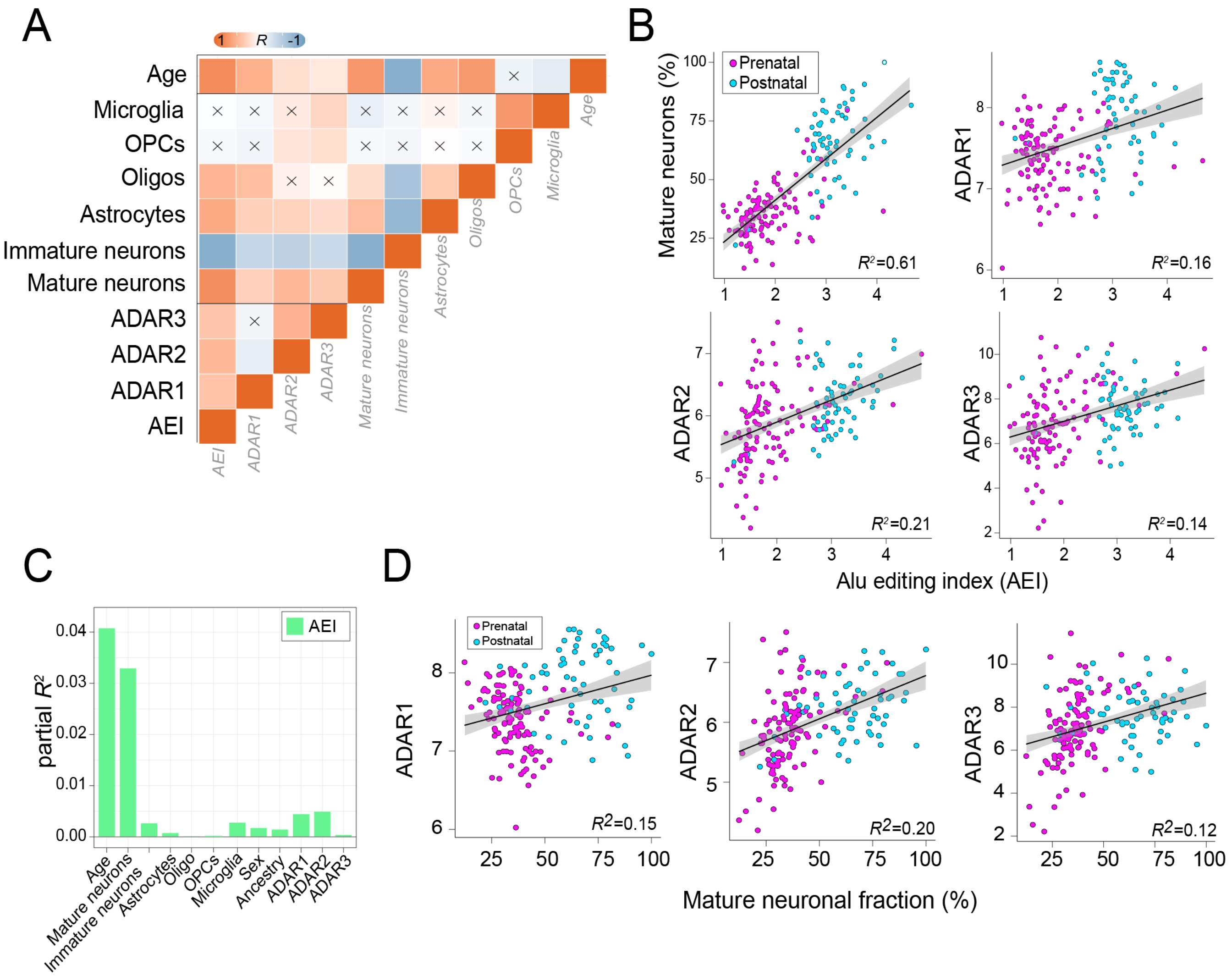


**Supplemental Figure 2. Correlations with the proportions of mature neurons.** (**A**) Pairwise pearson correlation coefficients measured all possible pairwise comparisons between chronological age, predicted cell type proportions (e.g. Microglia, Oligodendrocyte precursor cells (OPCs), Oligodendorcytes (Oligos), immature and mature neurons), ADAR expression and the Alu editing index. An X indicates a non-significant association. (**B**) Pairwise correlations between estimated mature neuronal cell type proportions and *ADAR1*, *ADAR2*, and *ADAR3*. (**C**) Partial R-squared computed for the AEI (y-axis) according to cell-type proportions and known technical and demographic factors. (**D**) Correlation (R-squared) by the AEI according to estimated mature neuronal proportions, *ADAR1*, *ADAR2*, and *ADAR3* expression (pink, prenatal; blue, postnatal samples).

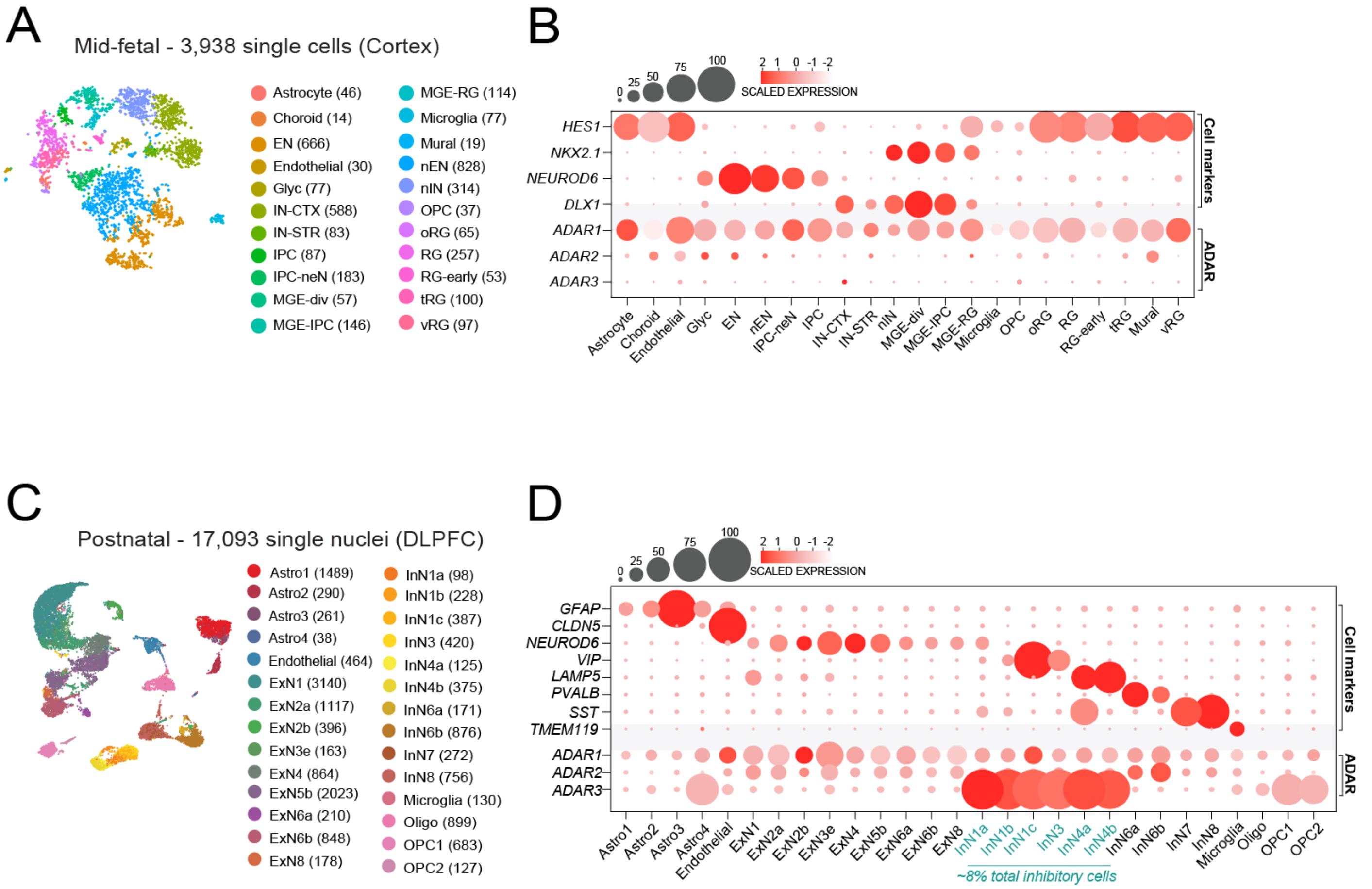


**Supplemental Figure 3. Single-cell RNA-seq analyses of *ADAR* expression.** (**A**) A total of 3,938 single cells were analyzed from human fetal cortical tissue and clustered using UMAP dimensionality reduction techniques. (**B**) A combination of early cell markers to define cell types and ADAR expression enzymes were plotted by detection rates per cell cluster (larger spheres indicate a larger percentage of the cells a given cluster expressing the gene). Here, ADAR2 and ADAR3 are lowly and sparsely expressed. (**C**) A total of 17,093 single nuclei were analyzed from human postnatal cortical tissue and clustered using UMAP dimensionality reduction techniques. (**D**) A combination of early cell markers to define cell types and *ADAR* expression enzymes were plotted by detection rates per cell cluster (larger spheres indicate a larger percentage of the cells a given cluster expressing the gene). Here, *ADAR2* and *ADAR3* are highly expressed and defined in specific cell subset classes. Notably *ADAR3* is expressed in a subclass of VIP and LAMP5 inhibitory cell types.

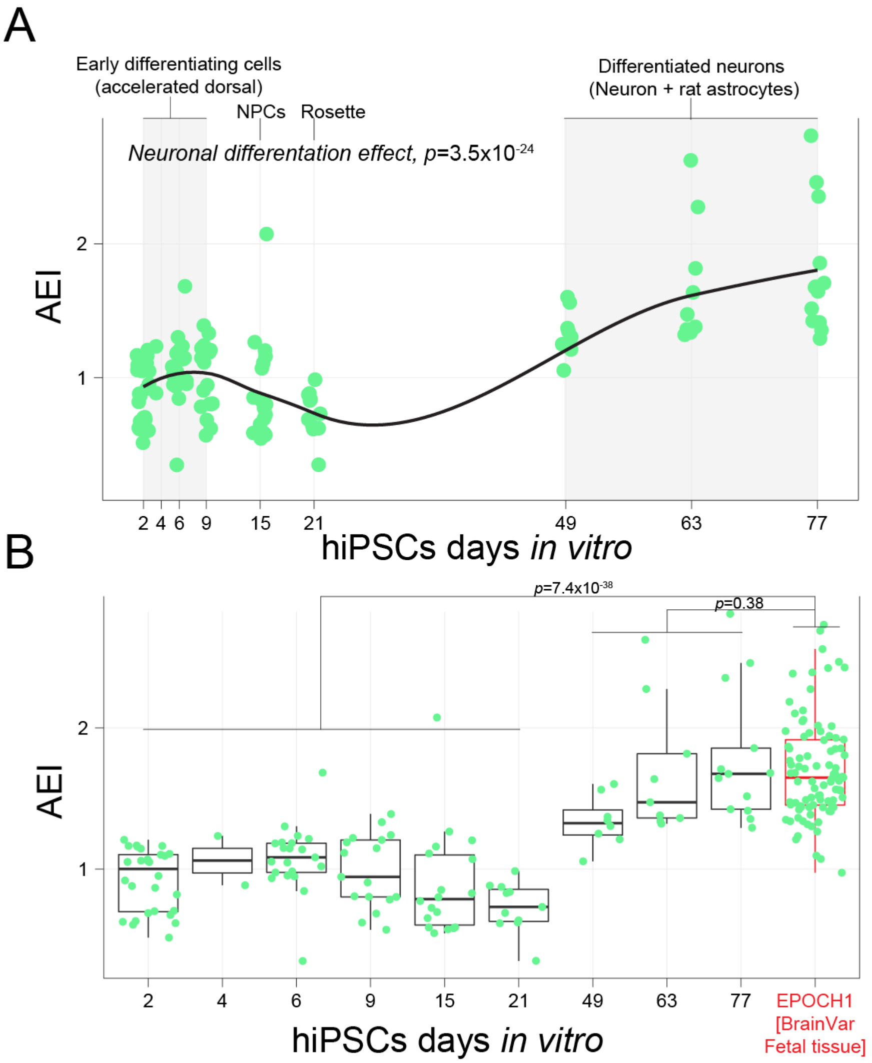


**Supplemental Figure 4. Alu editing index (AEI) in a hiPSC neuronal differentiation time course.** (**A**) AEI (y-axis) across 77 days in culture (x-axis) covering four main stages, early differentiating cells, neural progenitor cells, assembled neuroepithelial rosettes and more differentiated neuronal cell types. A loess curve was used to fit the data and a linear regression tested for significance with neuronal differentiation. (**B**) The AEI from early fetal DLPFC in BrainVar (Epoch 1, *n*=91) was compared to fully differentiated neurons and all remaining time points in culture. A linear regression tested for significance.

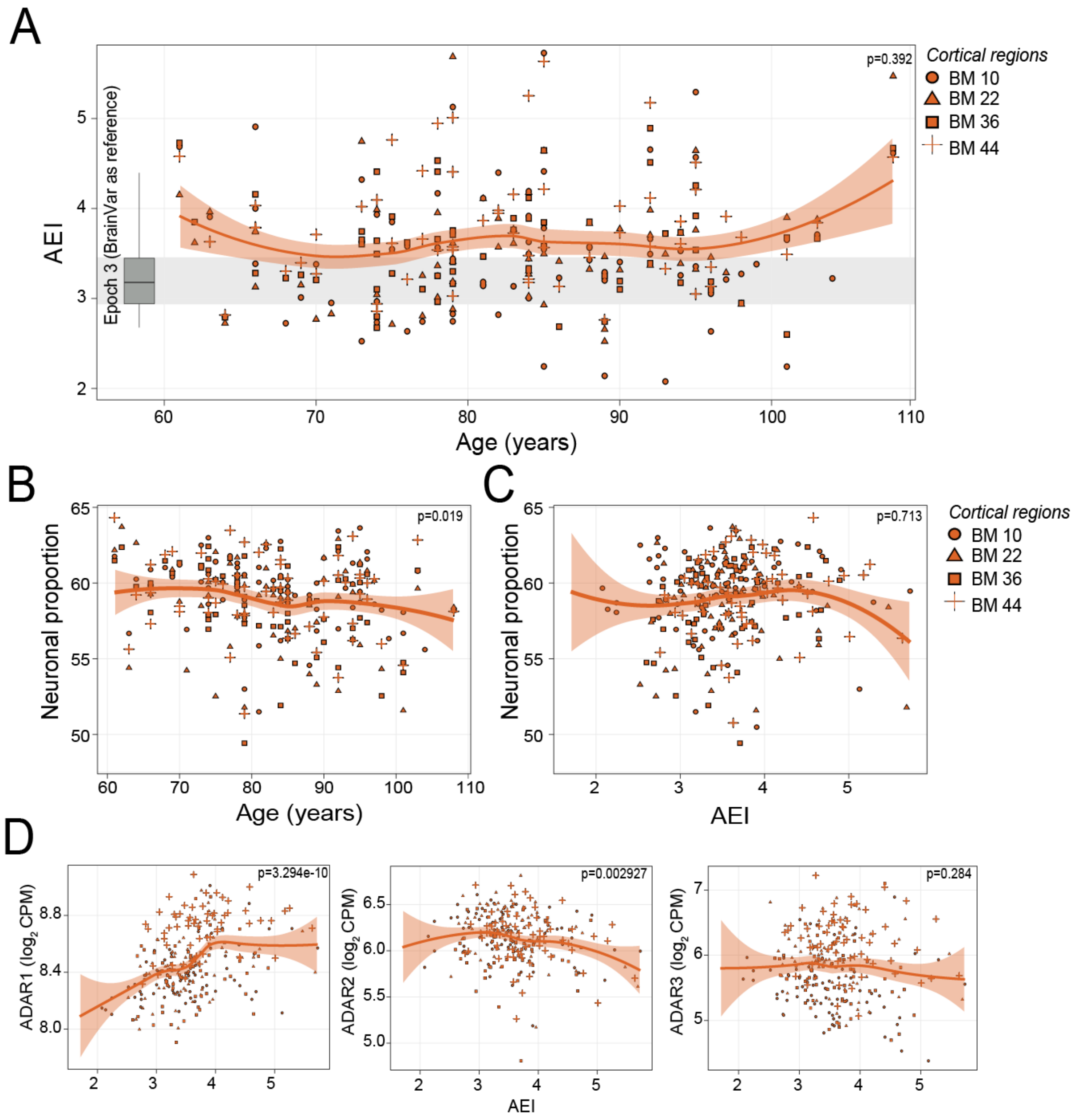


**Supplemental Figure 5. Alu editing index (AEI), neurons and ADARs in advanced age.** (**A**) AEI (y-axis) across four cortical areas throughout advanced age (years, x-axis). Inset boxplot and grey bar indicate AEI for the 3$^{rd}$ epoch of BrainVar. Association between estimated mature neuronal cell type proportions (y-axes) and (**B**) age in years (x-axis) and (**C**) the AEI (x-axis). (**D**) Association of AEI between *ADAR1*, *ADAR2* and *ADAR3* expression. For each test, a linear regression tested for significance. A loess curve was used to fit the data.

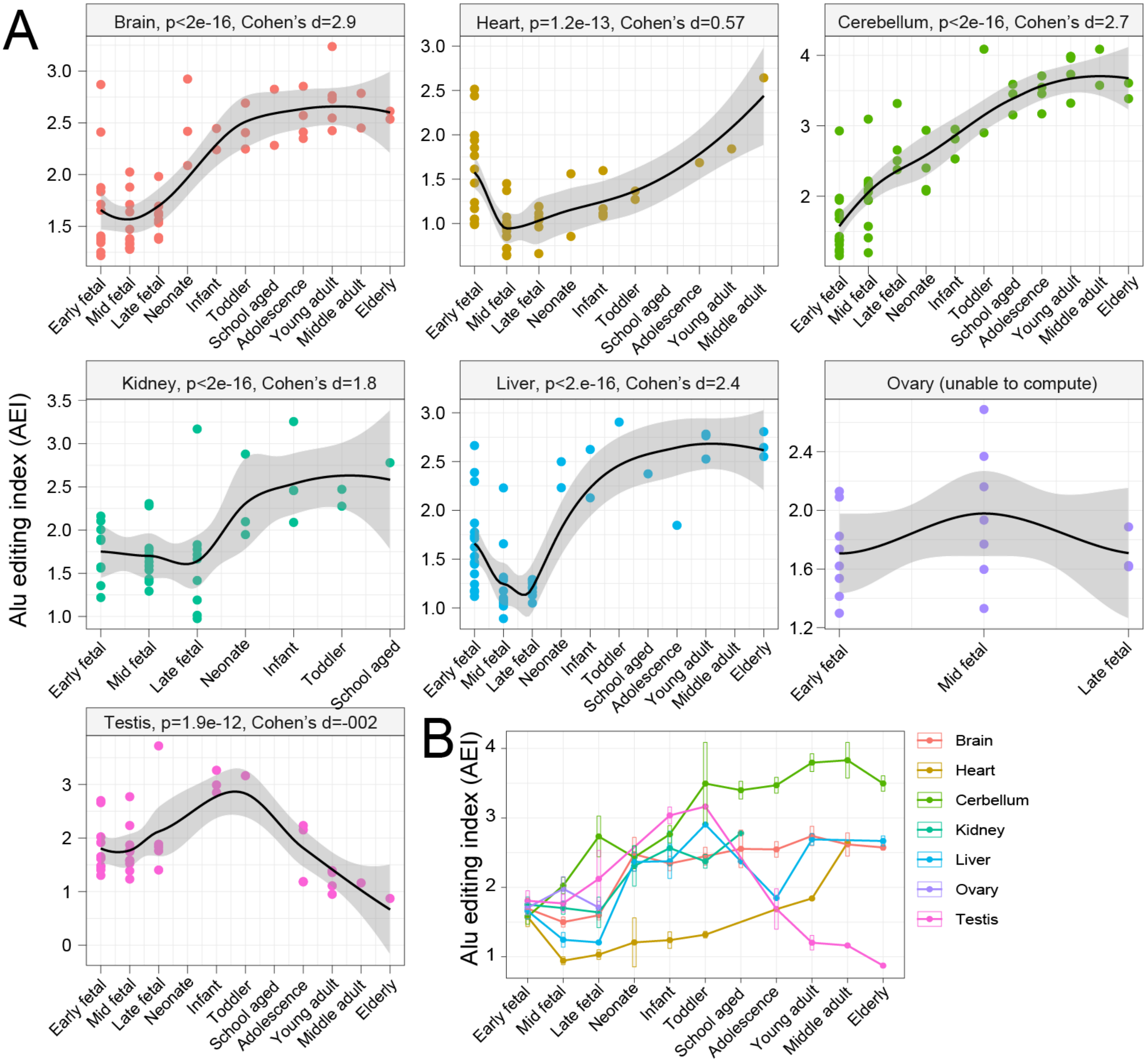


**Supplemental Figure 6. Alu editing index (AEI) of 6 human tissues across the lifespan.** (**A**) AEI (y-axis) across all stages of development (x-axis) for bulk brain (*n*=55), heart (*n*=48), cerebellum (*n*=55), kidney (*n*=40), liver (*n*=50) and testis (*n*=41). We also computed the AEI for ovary (*n*=18), which was sampled only during prenatal stages. For each test, a linear regression tested for significance. A loess curve was used to fit the data. (**B**) A comparative plot of the average AEI (and standard error) per time period for each tissue.

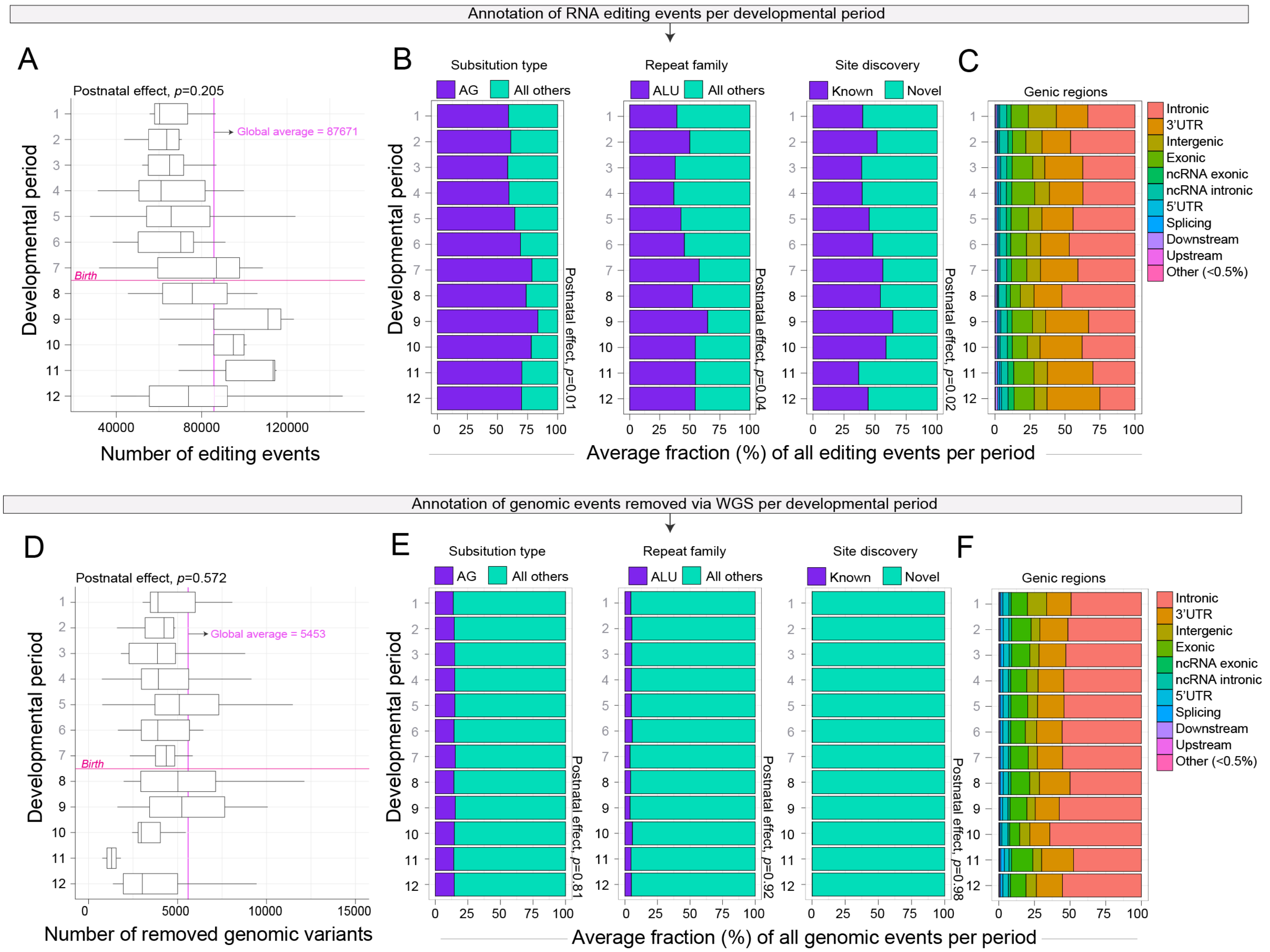


**Supplemental Figure 7. Annotation of true editing sites versus sites masked as genomic calls.** Annotation of bona fide RNA editing sites per developmental period: (**A**) The number of RNA editing sites (x-axis) detected per period (y-axis). The global average number of sites is marked with a pink line. (**B**) The percentage (%) of all unique sites per period partitioned by A-to-G substitutions, sites in Alu repeats and known sites catalogued in a RNA editing database. (**C**) The percentage of all unique editing sites per period according to the corresponding genic regions. Annotation of RNA editing sites that were removed following filtering by whole genome sequencing (WGS) data: (**D**) The number of RNA editing sites (x-axis) removed per period (y-axis). The global average number of RNA editing sites removed by paired WES is marked with a pink line (~5,453 sites). (**E**) The percentage (%) of all unique genomic sites per period partitioned by A-to-G substitutions, sites in Alu repeats and known sites catalogued in a RNA editing database. (**F**) The percentage of all unique genomic editing sites per period according to the corresponding genic regions. A linear regression was used to test each partition of the data across all developmental periods and corresponding P-values are displayed.

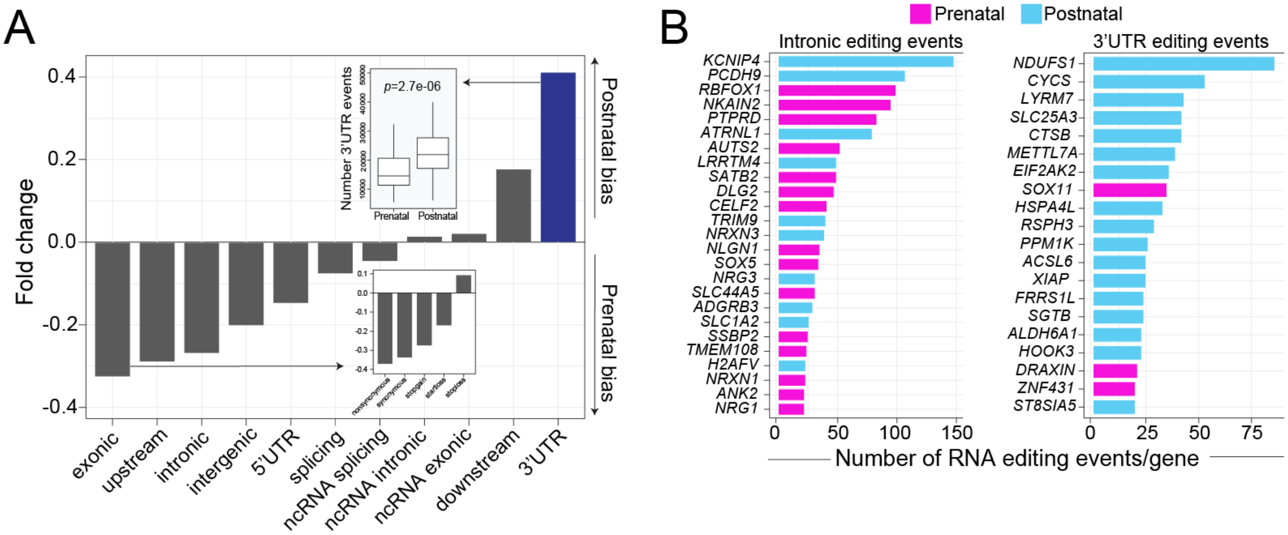


**Supplemental Figure 8. RNA editing in introns and 3'UTRs in prenatal- and postnatal-predominant sites.** (**A**) Fold-changes according to the total number of RNA editing sites catalogued by genic region illustrates a significant enrichment in the frequency of editing sites in 3'UTRs among sites specific to postnatal periods 8-12. A linear regression was used to test for an association between the frequency of events and developmental periods. (**B**) Genes that harbor a large number of selective RNA editing events in either intronic regions or 3'UTRs among prenatal and postnatal predominate sites. The top 20 genes were ranked and plotted for each category, respectively.

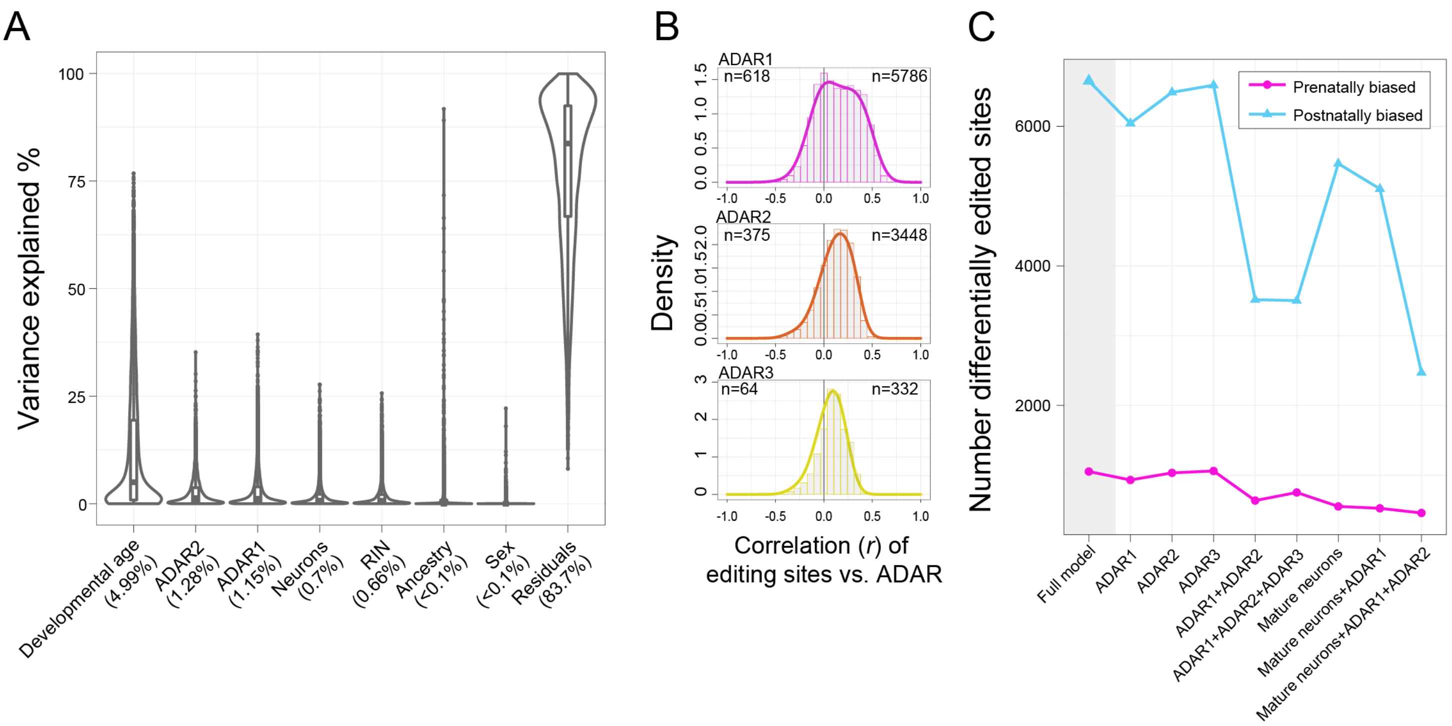


**Supplemental Figure 9. Variance in RNA editing rates explained.** (**A**) Linear mixed modelling was used to compute the percentage of RNA editing variance explained according to seven factors, which represent potential sources of variability. Differences in developmental (chronological) *ADAR2* and *ADAR1*, followed by mature neuron cell composition explains the largest amount of variability. (**B**) Pearson correlation coefficients for ADAR1, ADAR2 and ADAR3 as associated with 10,652 RNA editing sites. The total number of significant correlations that are either positively correlated (upper right corner) or negatively correlated (upper left corner) are displayed for each association. (**C**) The number of developmentally regulated sites (Adj. $P < 0.05$) following a combination of covarying for different factors. The full model adjusted for biological sex and ancestry. Subsequently, the contribution of ADAR1, ADAR2 and mature neuronal proportions were accounted for. The number of significant sites (y-axis) remaining after fitting each corresponding model (x-axis).

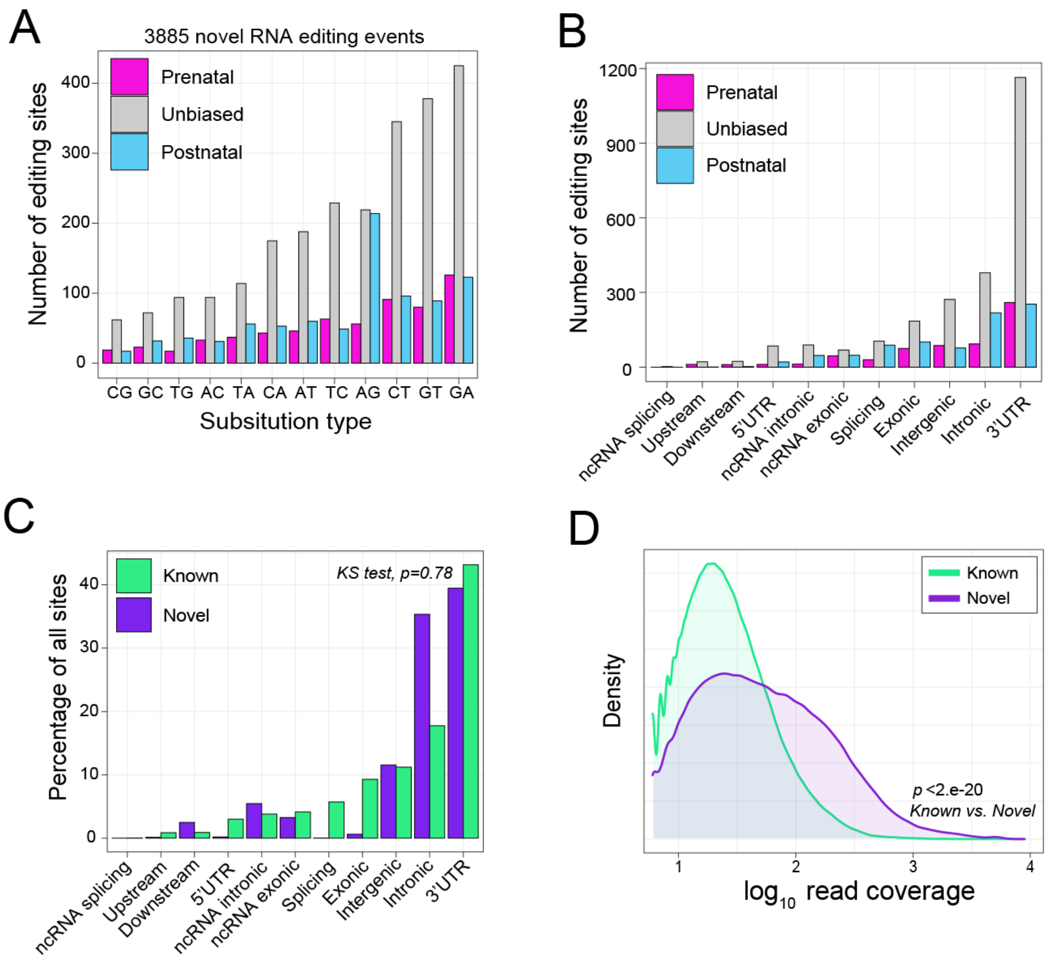


**Supplemental Figure 10. Annotation of putative novel RNA editing sites.** The current study focused predominately on A-to-G and C-to-T editing as enzymes have been described to catalyze these events. However, our *de novo* calling approach identified all possible types of editing, which we describe briefly here. A total of 3,885 putatively novel sites were identified. (**A**) The distribution of different substitution classes according to whether these sites were prenatal, postnatal or unbiased in either editing rates. (**B**) The distribution of different genic regions according to whether these sites were prenatal, postnatal or unbiased in either editing rates. (**C**) Overall, the distribution across genic regions was no different from known sites. Significance was tested using a Kolmogorov-Smirnov test ($p$=0.78). (**D**) Novel sites also displayed significantly more read coverage ($\log_{10}$ scale, x-axis) relative to known RNA editing sites. Significance was tested using a Mann-Whitney U test.

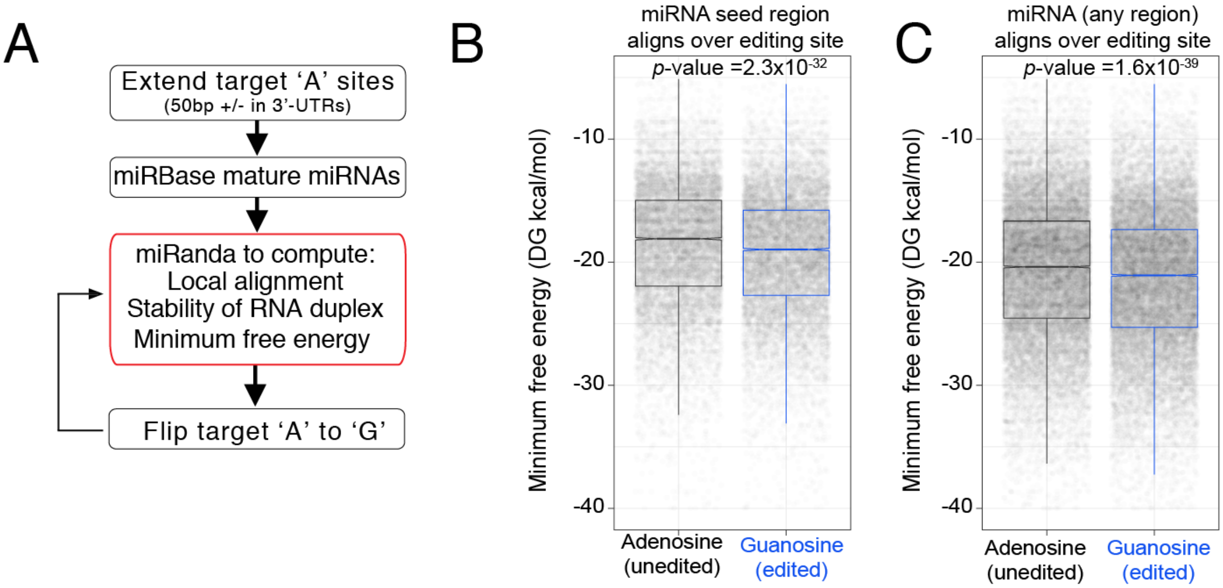


**Supplemental Figure 11. miRNA binding energy.** (**A**) Work-flow used to compute differences of miRNA minimum free energy on edited and un-edited 3'UTRs. Differences in miRNA minimum free energy computed for only high confident local alignments for (**B**) miRNA seed regions or (**C**) any region of a miRNA that overlap with an editing site in a 3'UTR. Significance was tested using a Mann-Whitney U test.

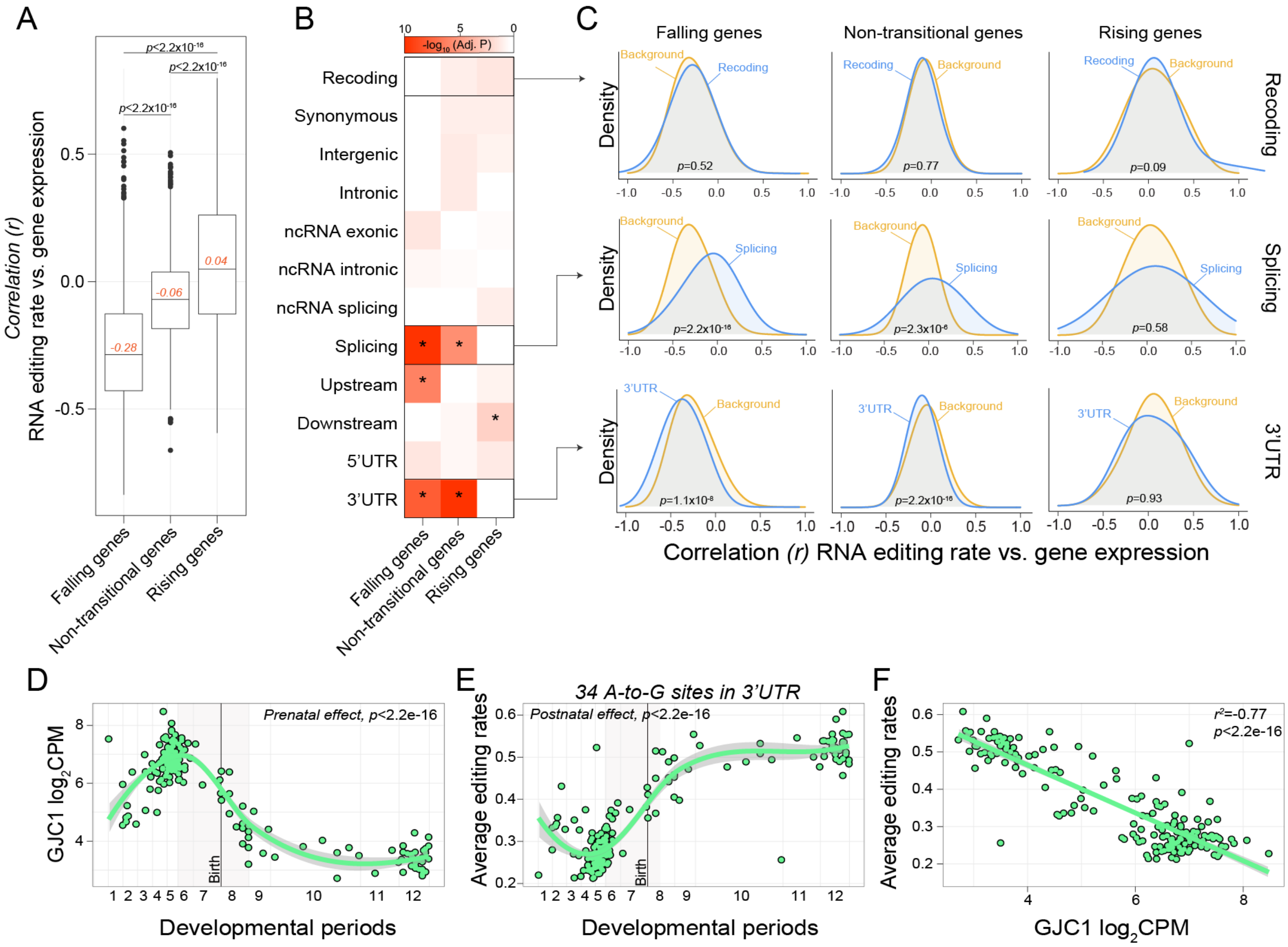


**Supplemental Figure 12. Correlation between RNA editing sites and corresponding gene expression profiles.** (**A**) RNA editing rates were correlated to the corresponding gene expression using a Pearson's correlation coefficient. We anchored this analysis around developmental trajectories of genes catalogued into three groups: falling, non-transitional or rising genes across development. (**B**) Subsequently, for each group, the correlation between RNA editing sites versus corresponding gene expression profiles were parsed based on the genic location of the editing event. Each distribution was compared to a background distribution (all other RNA editing sites vs. gene expression associations) and tested for significance using a Mann-Whitney U test. Here asterisks indicate p-value < 0.05. (**C**) A visual of the distributions of editing sites vs. gene expression correlations (x-axis) for sites that map to exons are recode amino acids, splice site regions and introns. (D-F) Gene GJC1 is an example of a gene that is prenatally biased in expression, but has 34 A-to-G editing events in the 3'UTR that all increase in editing efficiency over development and are negatively correlated (i.e. more editing in 3'UTR reflects reduced expression).

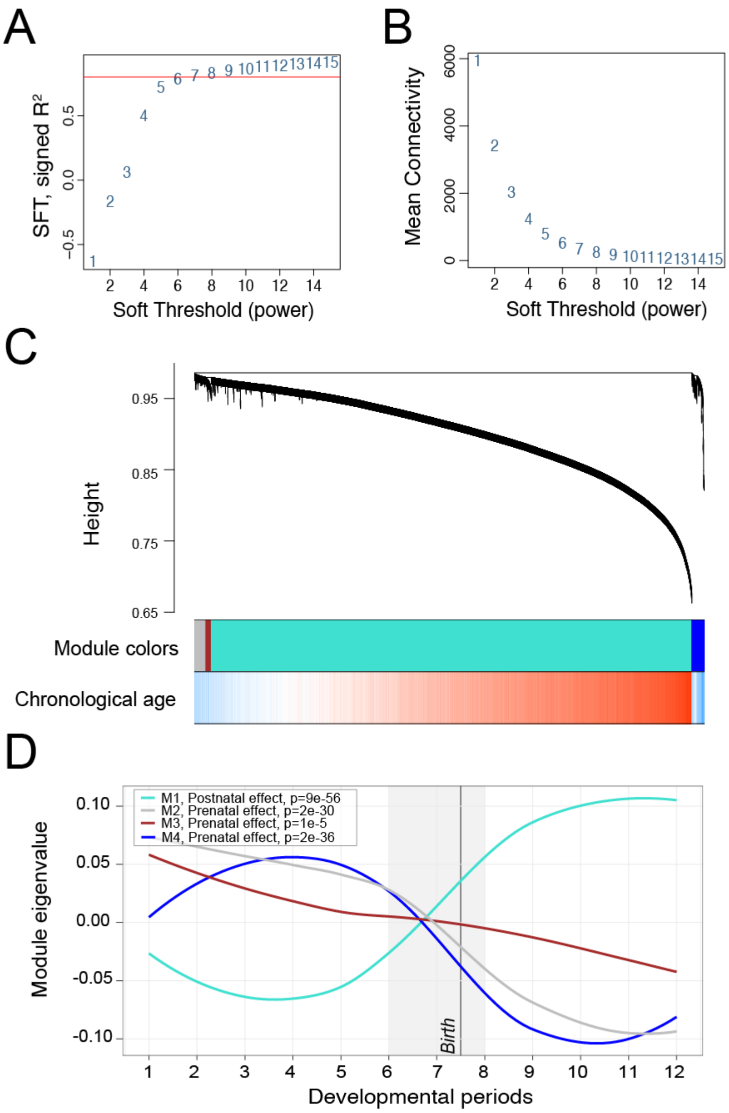


**Supplemental Figure 13. Weighted correlation network analysis** (**A**) To determine the optimal soft threshold (beta-value) (y-axis) to weigh the network, we evaluated a measure of scale free topology (SFT, y-axis) relative to a series of beta-values. Here, we used a beta-value of 7 to construct our network. (**B**) Similarly, network mean connectivity (y-axis) was compared across a series of beta-values. (**C**) All sites were used to construct a weighted network to identify discrete groups of co-regulated sites (modules) that could be functionally interrogated. A hierarchical cluster tree (dendrogram) is displayed. Each line represents an editing site (leaf) and each low-hanging cluster represents a group of co-edited sites with similar network connections (branch) on the tree. The first band underneath the tree indicates the four detected, and subsequently analyzed, network modules. The second band indicates correlation with chronological age, where red lines indicate positive editing site-age relationships and blue lines indicate negative editing site-age relationships. (**D**) The editing variation in each module was summarized to a module eigengene (ME) value (y-axis), equivalent to the first principal component and plotted across all developmental periods (x-axis). Linear regression analyses tested the associations between all module MEs and development.

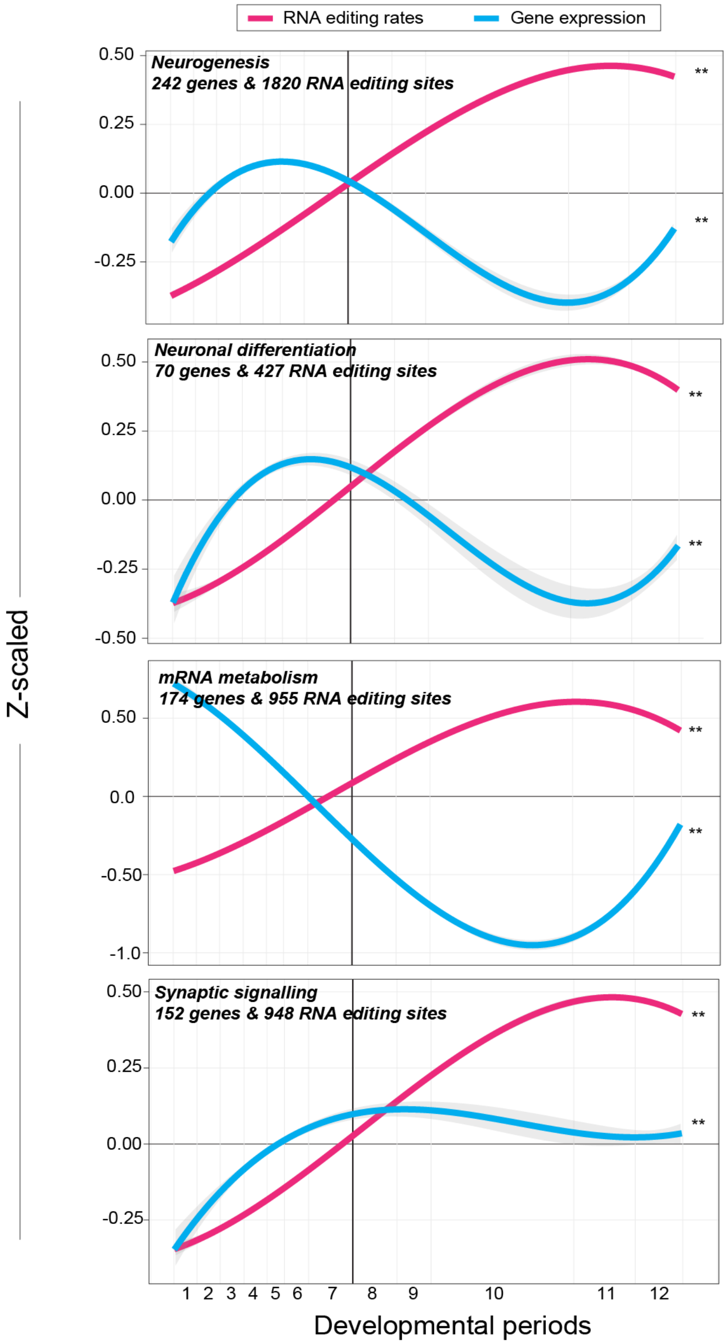


**Supplemental Figure 14. Paired RNA editing and gene expression for candidate gene sets.** Paired RNA editing rates and gene expression profiles for gene sets known to peak in expression during prenatal periods (neurogenesis, neuronal differentiation, mRNA metabolism) and one gene set known to peak in expression during postnatal periods (synaptic signaling). All measurements are Z-scaled for comparison. Linear regression analyses were used to test for significant association with age and ** indicates associations with *P*-value < $2.6 \times 10^{-16}$.

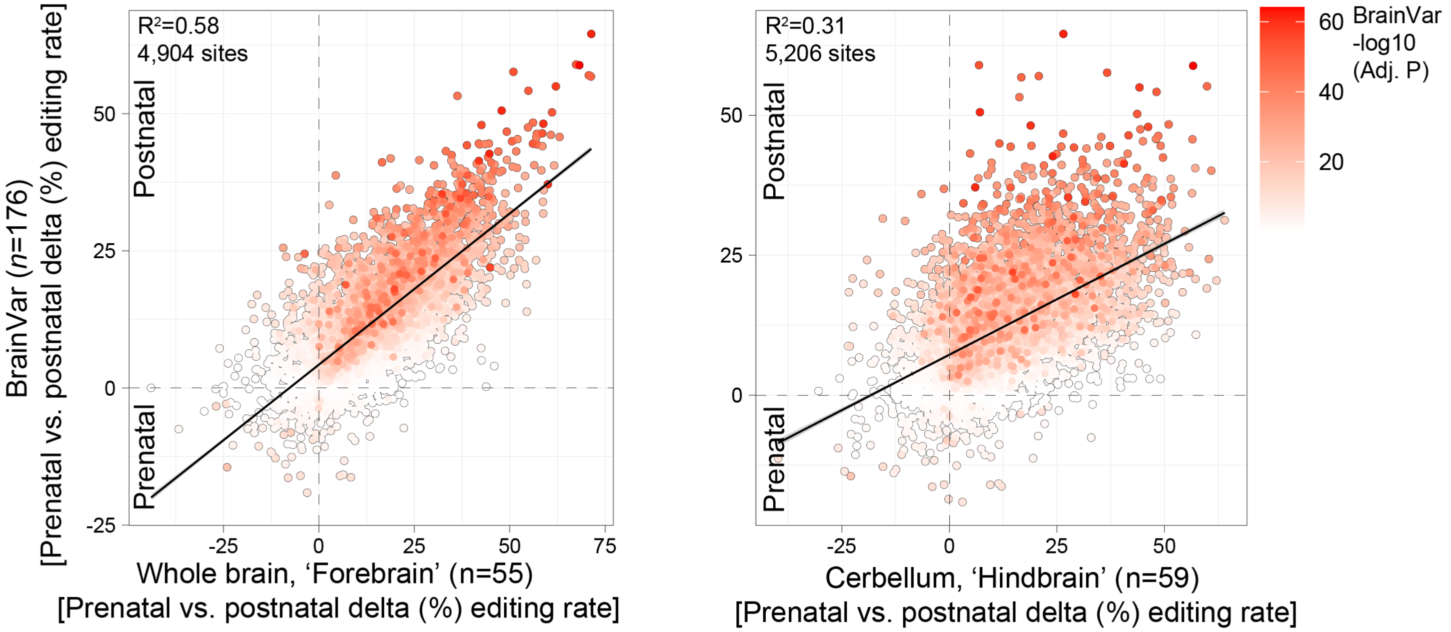


**Supplemental Figure 15. Validation of developmentally regulated selective editing sites.** The developmental change (delta) in selective editing rates from BrainVar (y-axes) are compared to those in a smaller independent developmental data set of forebrain (x-axis, left) and cerebellar (x-axis, right) tissues. Moderated t-tests in the limma package were used to test the developmental effect in these independent data, and a linear regression was used to test the transcriptome-wide concordance of delta editing rates. The number of consistently detected and validated sites are displayed for each analysis. Points are shaded by –log10 adjusted p-value based on the BrainVar analysis.

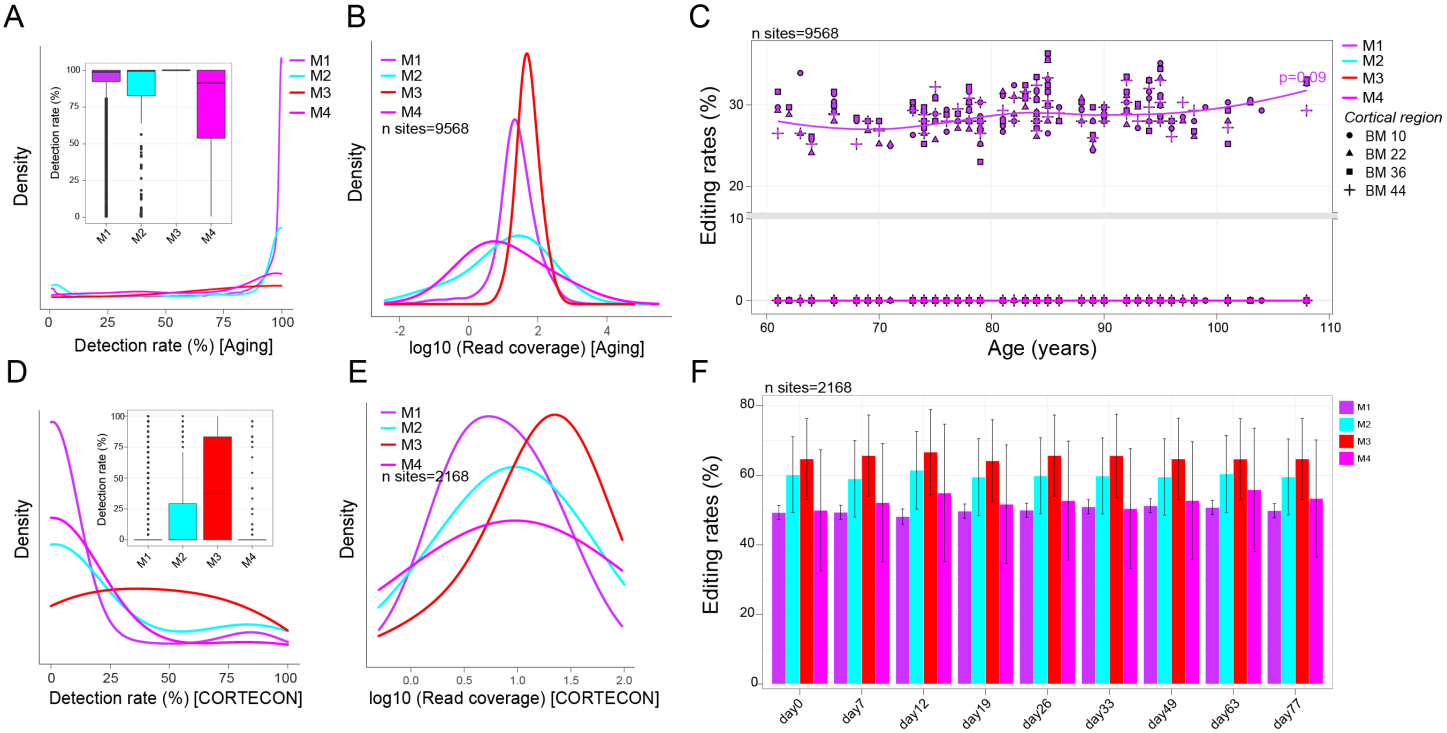


**Supplemental Figure 16. Validation of modules of coordinated editing sites.** For 261 cortical samples during advanced age: (**A**) Detection rates (%) of all modules defined by at least 10 reads covering each site in a module across all 261 samples. Inset boxplots reflect the average detection rates for all modules. (**B**) For sites with at least 70% detection rate (N=9568 sites), $log_{10}$ read coverage was queried all sites for each module. (**C**) Editing rates (y-axis) for all sites (N=9568 sites) across all four modules were displayed across advanced aging (x-axis). Significance was determined using a linear regression model. For 24 hESC samples during *in vitro* corticogenesis: (**D**) Detection rates (%) of all modules defined by at least 10 reads covering each site in a module across all 24 samples. Inset boxplots reflect the average detection rates for all modules. (**E**) For sites with at least 70% detection rate (N=2168 sites), $log_{10}$ read coverage was queried all sites for each module. (**F**) Editing rates (y-axis) for all sites (N=2168 sites) across all four modules were binned by days post neuronal induction (x-axis).

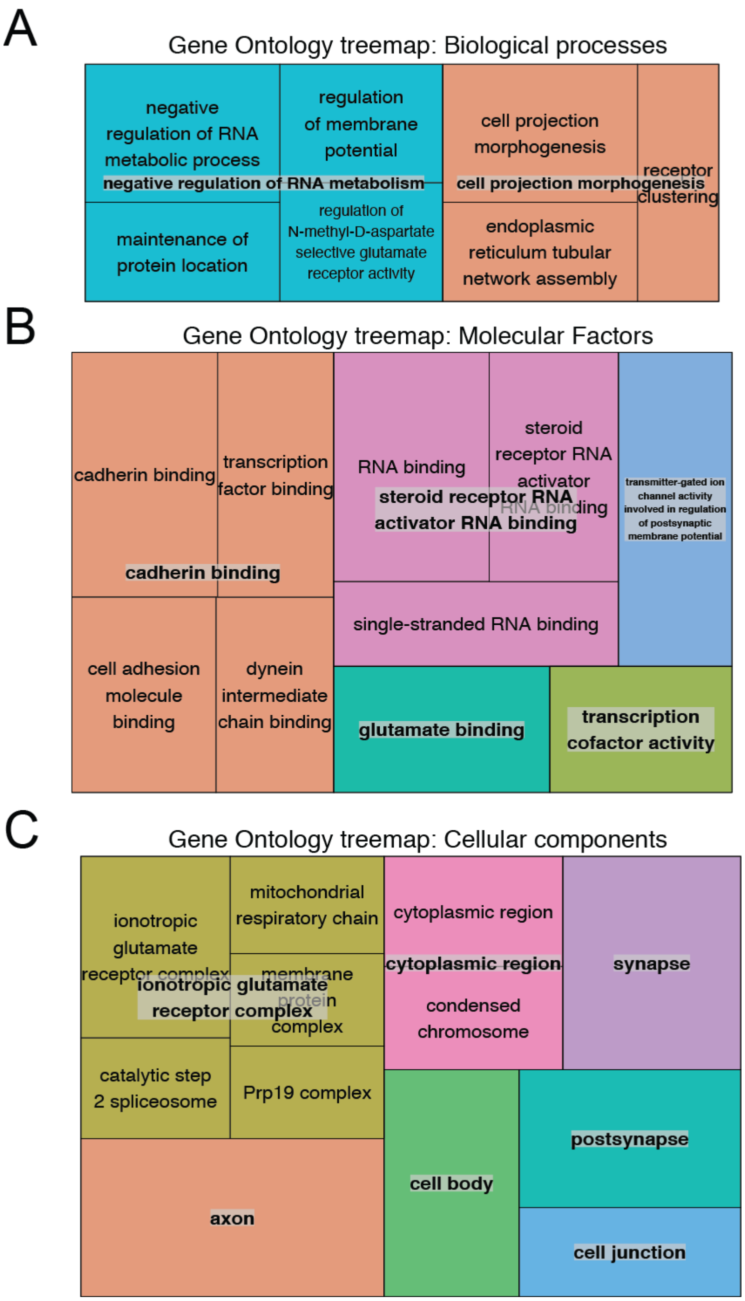


**Supplemental Figure 17. Semantic reduction of ontologies related to RNA recoding sites.** REVIGO summarizes extended lists of gene ontology (GO) terms using semantic similarity measures. ToppFun gene set enrichment analysis yielded several GO terms with a Benjamini-Hochberg adjusted *P*-value < 0.05, which were used as input to REVIGO. This analysis condensed significantly enriched gene sets specific to developmentally regulated RNA recoding sites according to (**A**) GO biological processes, (**B**) GO molecular functions and (**C**) GO cellular components.

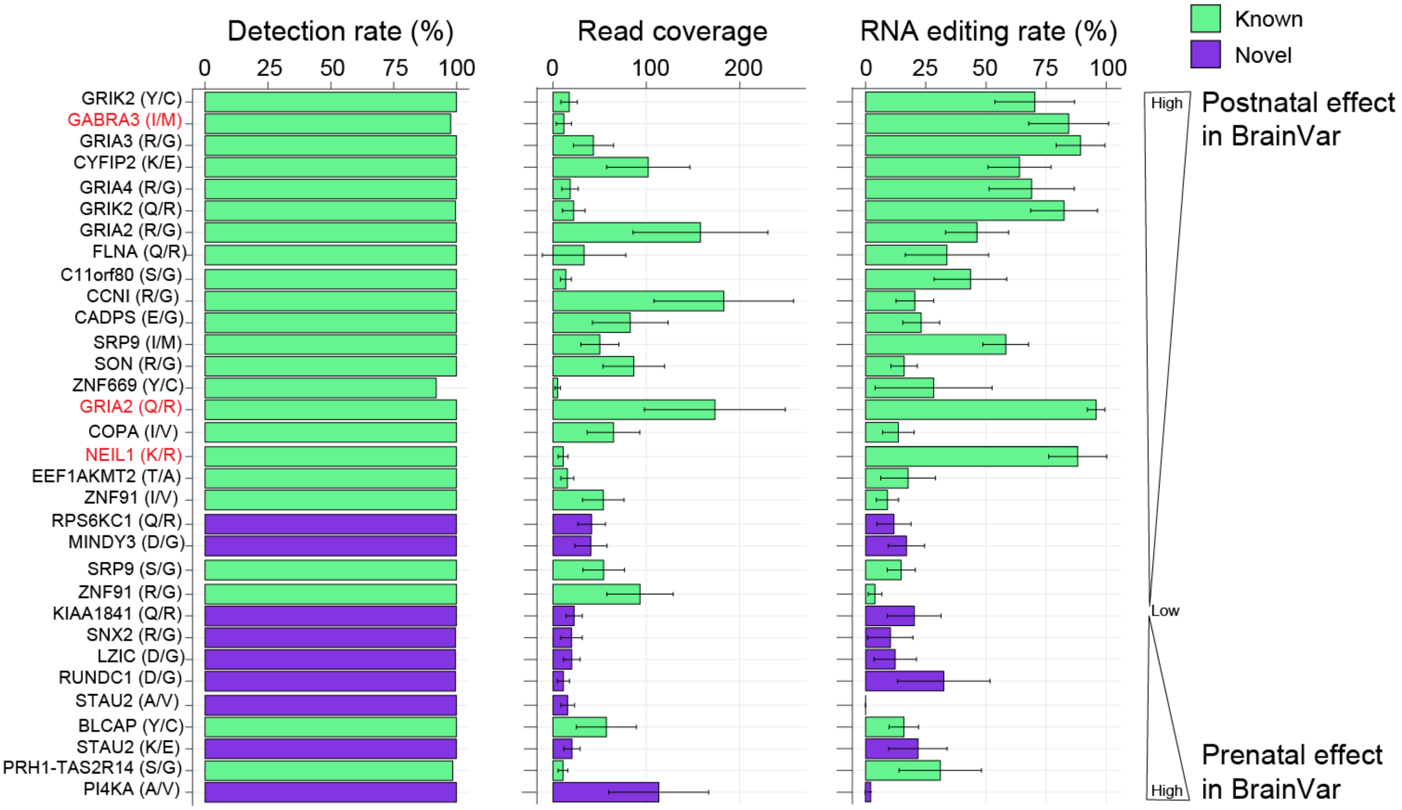


**Supplemental Figure 18. Validation of temporal recoding sites in advanced age.** A total of 32 recoding sites were identified with significant changes in editing efficiencies through development. Here we query these sites in advanced age across 261 cortical samples. (**A**) Detection rates (%), (**B**) read coverage, and (**C**) RNA editing rates for each site across 261 independent cortical samples.

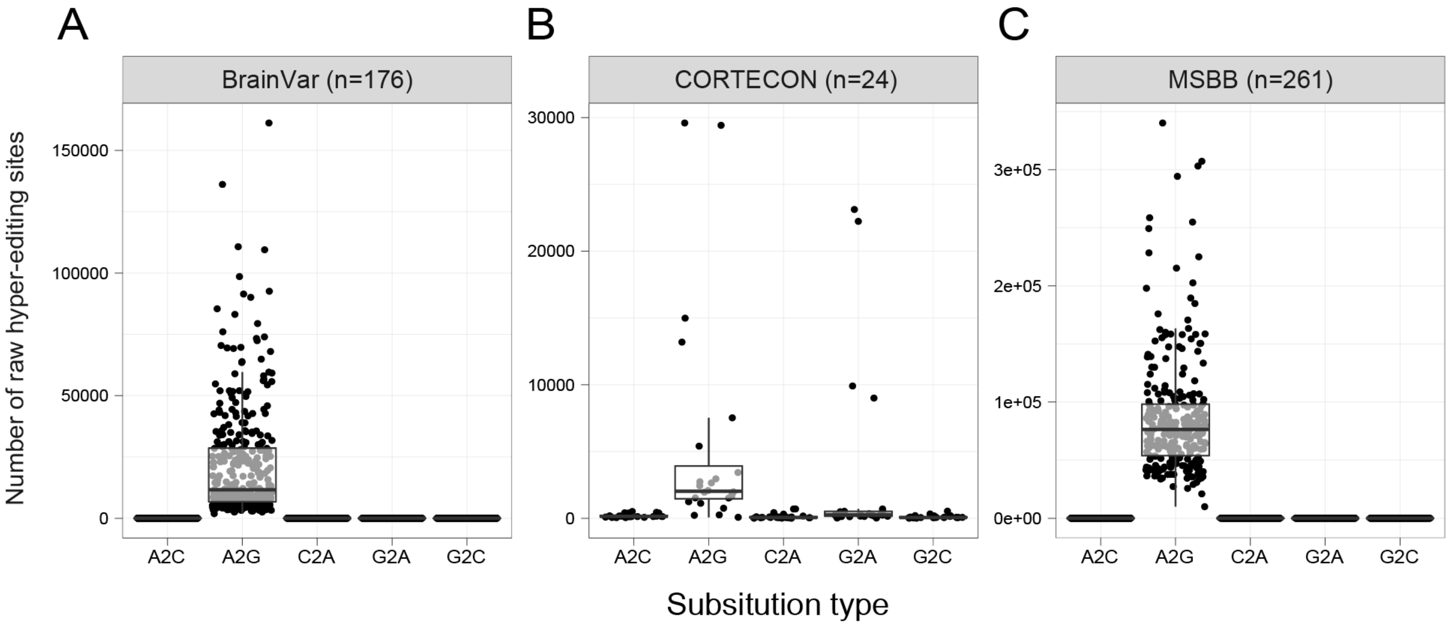


**Supplemental Figure 19. A massive A-to-G RNA hyper-editing signal among raw calls.** We queried a five different hyper-editing substitution types and observed a massive A-to-G signal among raw calls (prior to filtering further annotations and filtering of genomic sites and other artifacts, see Methods) for (**A**) BrainVar samples, (**B**) CORTECON samples and (C) MSBB samples. Thus, our downstream analyses focused explicitly on A-to-G hyper editing events.

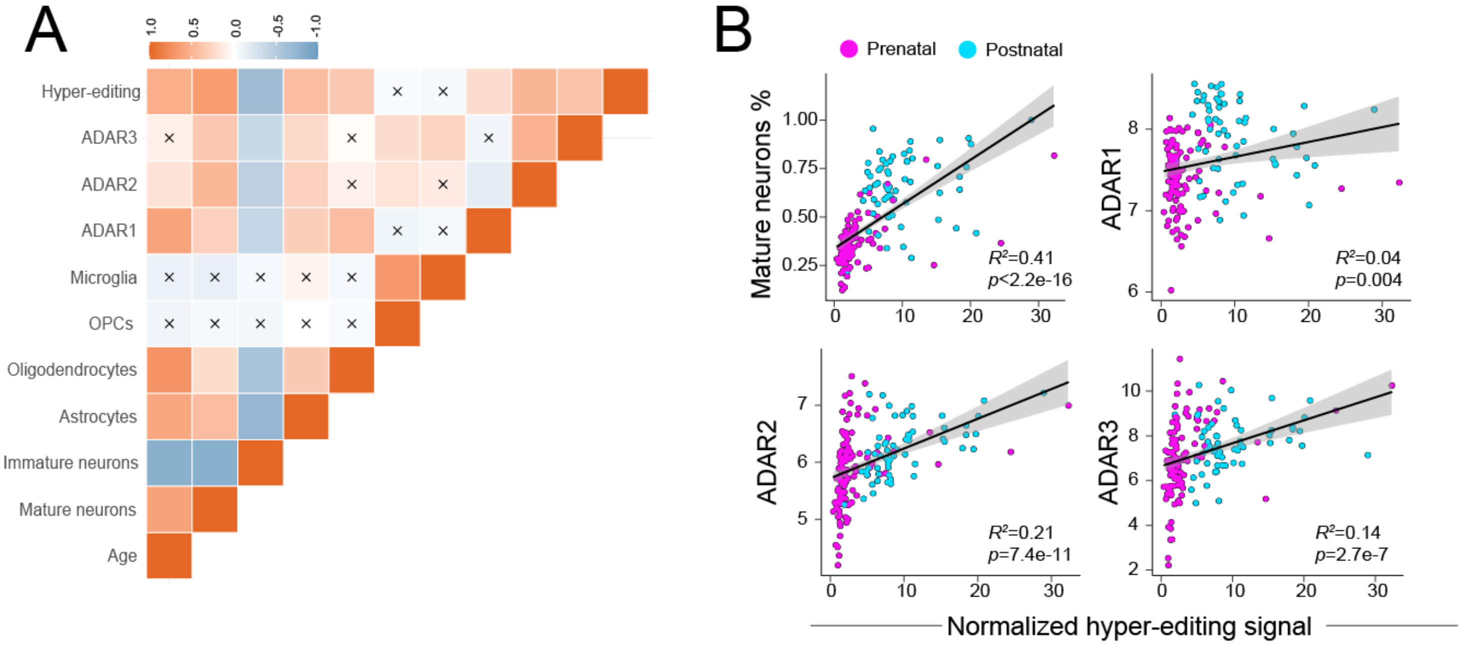


**Supplemental Figure 20. Variance explained in the A-to-G hyper-editing signal.** (**A**) Pairwise Pearson's correlations computed associations between the number of filtered high-confidence A-to-G hyper-editing sites with ADAR expression, estimated cell types and chronological age. (**B**) Pairwise linear regression analyses between the normalized A-to-G hyper-editing signal and the estimated proportion of mature neurons, *ADAR1*, *ADAR2* and *ADAR3*.

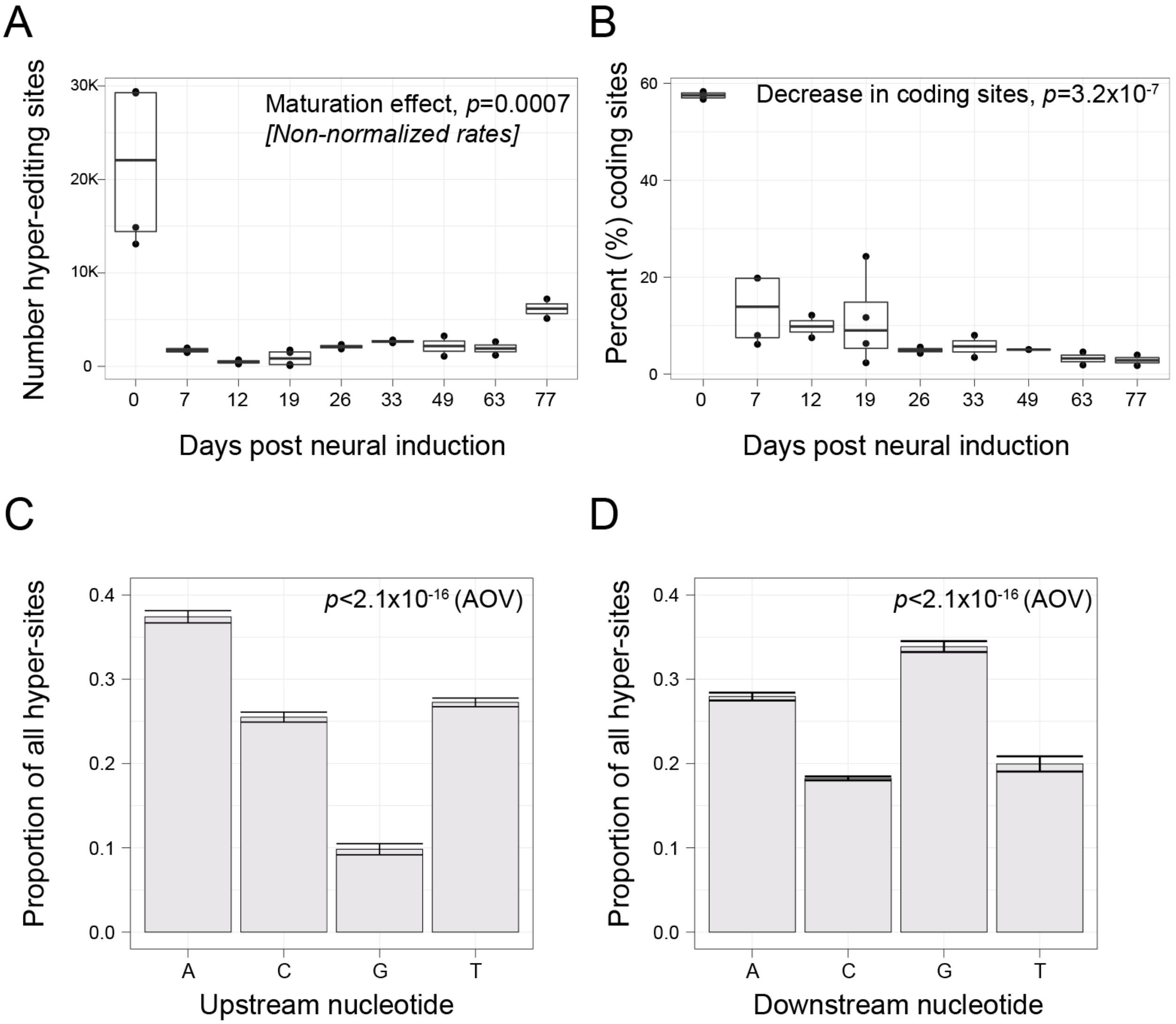


**Supplemental Figure 21. A-to-G hyper editing in an *in vitro* model of corticogenesis.** (**A**) The total number of high-quality A-to-G hyper-editing sites (y-axis) across days post neuronal induction (x-axis). A linear regression tested the number of sites across 77 days in culture. Note that day 0 indicates pluripotency and was dropped from this analysis as it was an extreme outlier in A-to-G hyper editing signal. (**B**) The majority of A-to-G hyper-editing signal during pluripotency occurred in coding regions, which are commonly rare events. Local sequence motifs for all high-confidence A-to-G hyper editing sites indicate a (**C**) depletion of guanosine 1bp upstream and (**D**) an enrichment of guanosine 1bp downstream the target adenosine. Analysis of variance was used to test for significance.

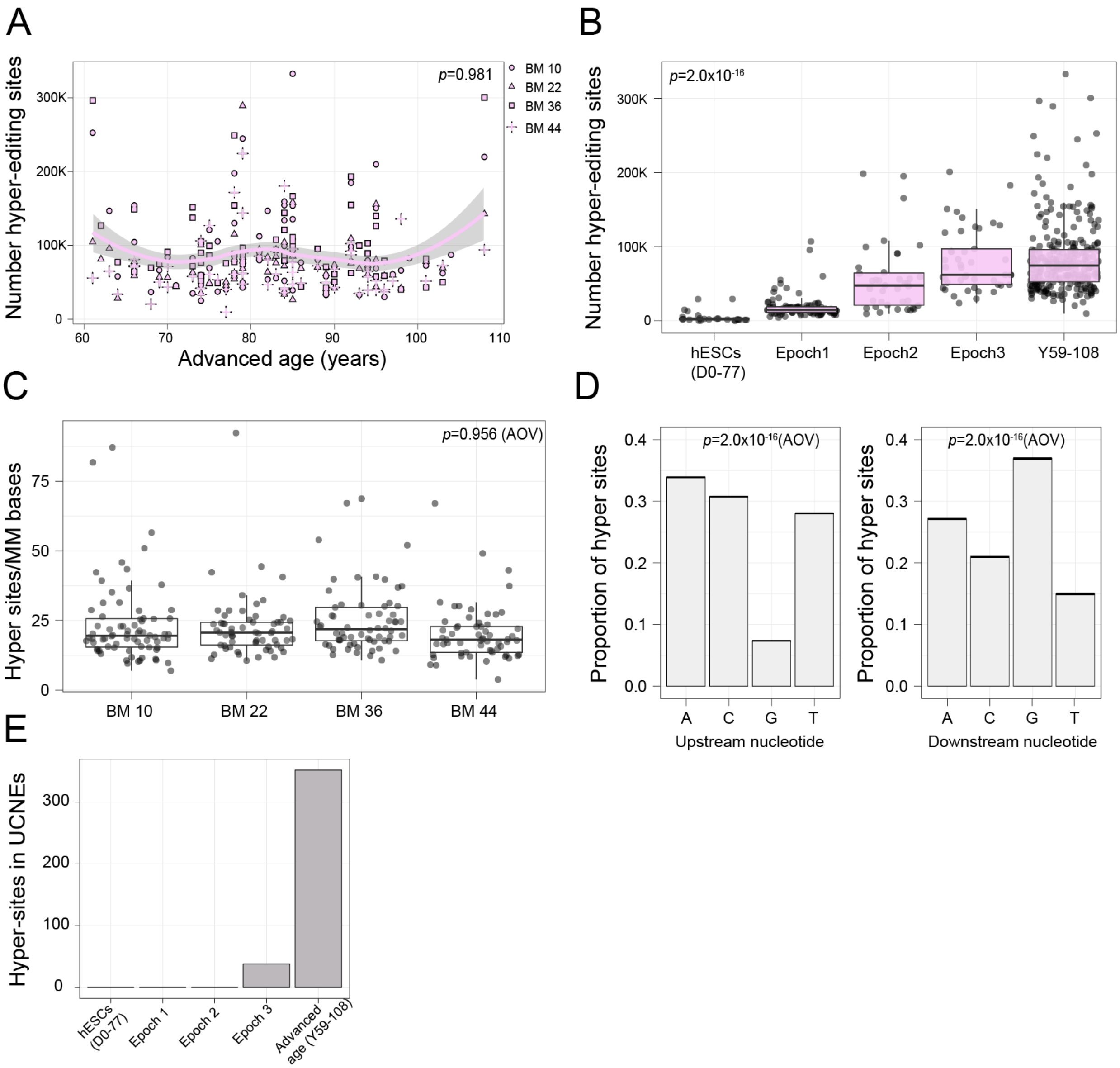


**Supplemental Figure 22. A-to-G hyper editing in during advanced age.** (**A**) The total number of high-quality A-to-G hyper-editing sites (y-axis) across advanced age (x-axis). A linear regression tested the number of sites across 58 years to 108 years of age. (**B**) The total number of high-quality A-to-G hyper-editing sites (y-axis) across all samples, including hESCs (n=24), epoch 1, epoch 2 and epoch 3 (BrainVar), and all advanced age samples (x-axis). A linear regression was used to test for significance. (**C**) Normalized hyper-editing rates display consistent A-to-G hyper-editing signal across four cortical areas during advanced age. (**D**) Local sequence motifs for all high-confidence A-to-G hyper editing sites indicate a depletion of guanosine 1bp upstream and an enrichment of guanosine 1bp downstream the target adenosine. Analysis of variance was used to test for significance. (**E**) The number of A-to-G hyper-editing sites that reside in ultra-conserved non-coding elements (UCNEs) of the genome across hESCs, epoch 1, epoch 2 and epoch 3 (BrainVar), and all advanced age samples (x-axis).

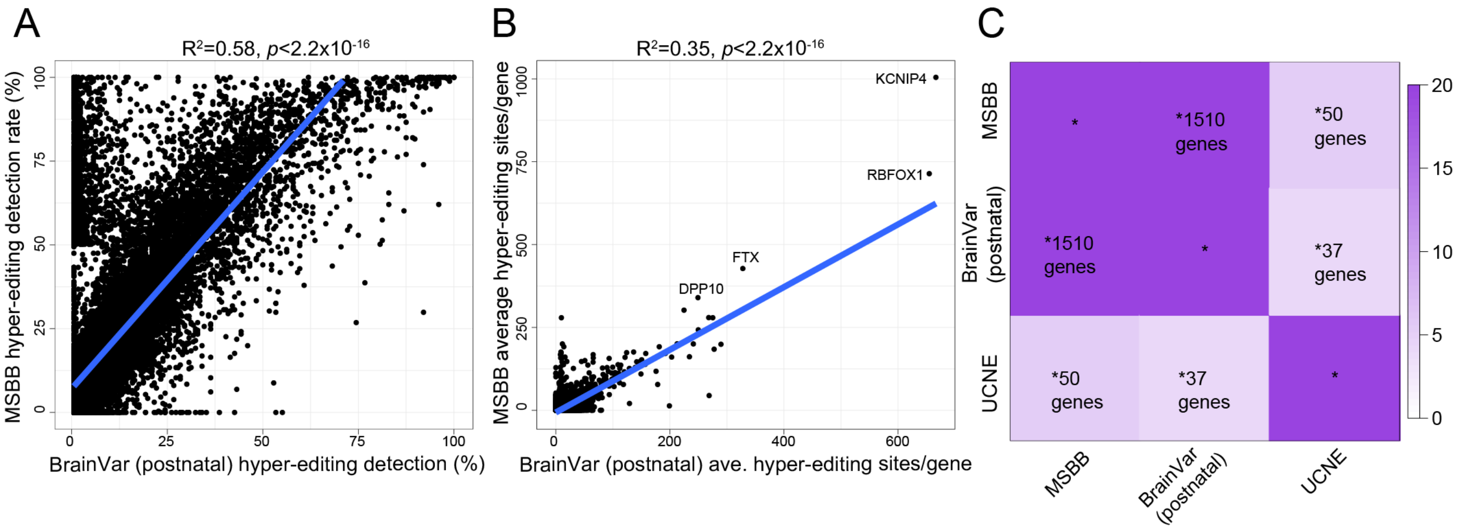


**Supplemental Figure 23. Validation of genes enriched with A-to-G hyper editing in advanced age. (A)** Detection rates (%) of a hyper-editing sites per gene across all samples under study. Here postnatal BrainVar samples (periods 8-12) (x-axis) and all advanced aging (MSBB) samples (y-axis) were used. Overall, the frequency of gene-specific hyper-editing was consistent across the two independent cohorts ($R^2$=0.58). (**B**) The average number of hyper-editing sites per gene across all BrainVar samples (x-axis) and all advanced aging samples (y-axis), illustrate a shift for increased hyper-editing signal during advanced age. (**C**) Over-representation analysis of genes with detectable hyper-editing in at least 40% of the entire cohort under-study. We used this as an unsupervised measure as the hyper-editing signal was enriched postnatally and it captured a broader, yet still specific, developmental window. We find an enrichment of hyper-edited genes between postnatal BrainVar samples and advanced age (MSBB), between advanced age and genes harboring ultra-conserved non-coding elements (UCNEs), and between postnatal BrainVar samples and UCNEs. A Fisher's exact test was used to test for significance.

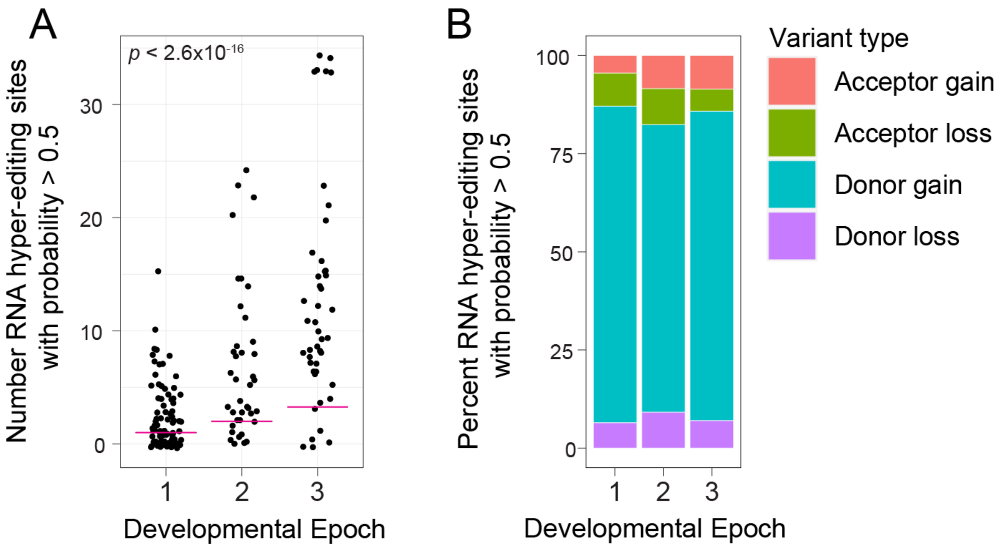


**Supplemental Figure 24. SpliceAI predictions of splice altering intronic RNA hyper-editing sites. (A)** The total number of splice altering intronic RNA hyper-editing events (y-axis) per donor according to developmental epoch (x-axis). We used a cut-off of $\geq$ 0.5 as a probability to interpret whether an editing site is splice altering. A linear model tested the association between epoch and number of sites with probability $\geq$0.5. (**B**) The percentage of all sites with probability $\geq$0.5 according to their variant type, whereby the majority of splice altering variants were predicted to be donor gain events.

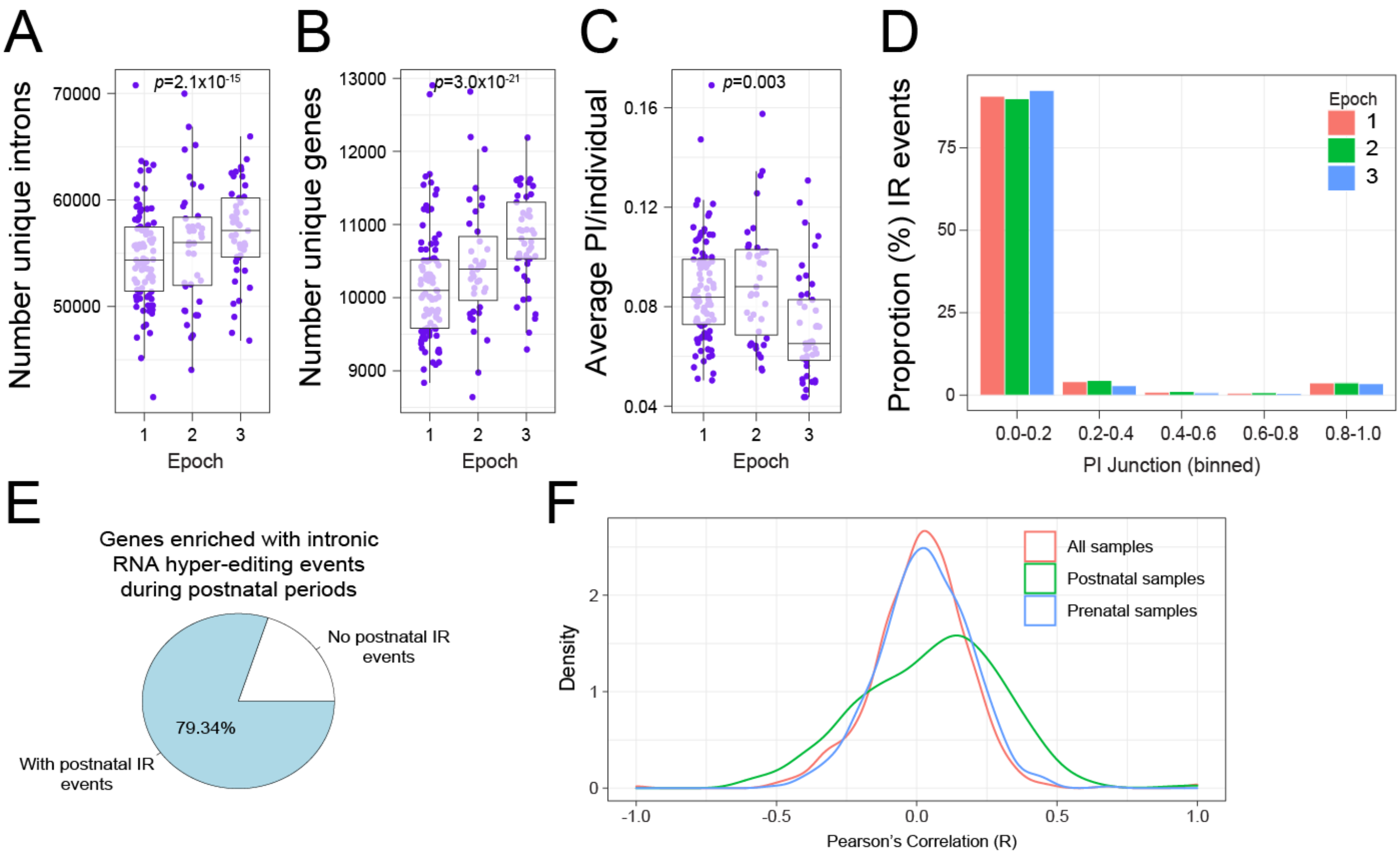


**Supplemental Figure 25. Intron retention events across cortical development.** The total number of (**A**) unique introns and (**B**) unique corresponding genes with detectable IR. We specifically focused on introns do not partly overlap with exons or with other introns. (**C**) Percent of introns (PI) was averaged across IR events per donor and is defined by inclusion counts divided by the sum of inclusion and skipping junction counts. Analysis of variance was used to test for significance across all three developmental epochs. (**D**) Frequency of PI metrics binned by epoch. (**E**) The fraction of genes enriched with intronic RNA hyper-editing sites during postnatal periods that were also associated with elevated IR. (**F**) Pearson's correlation coefficient of PI metric vs paired gene expression for 583 postnatally hyper-edited genes with measurable IR events. We used the top PI value per gene to compute association with gene expression ($log_2$CPM).

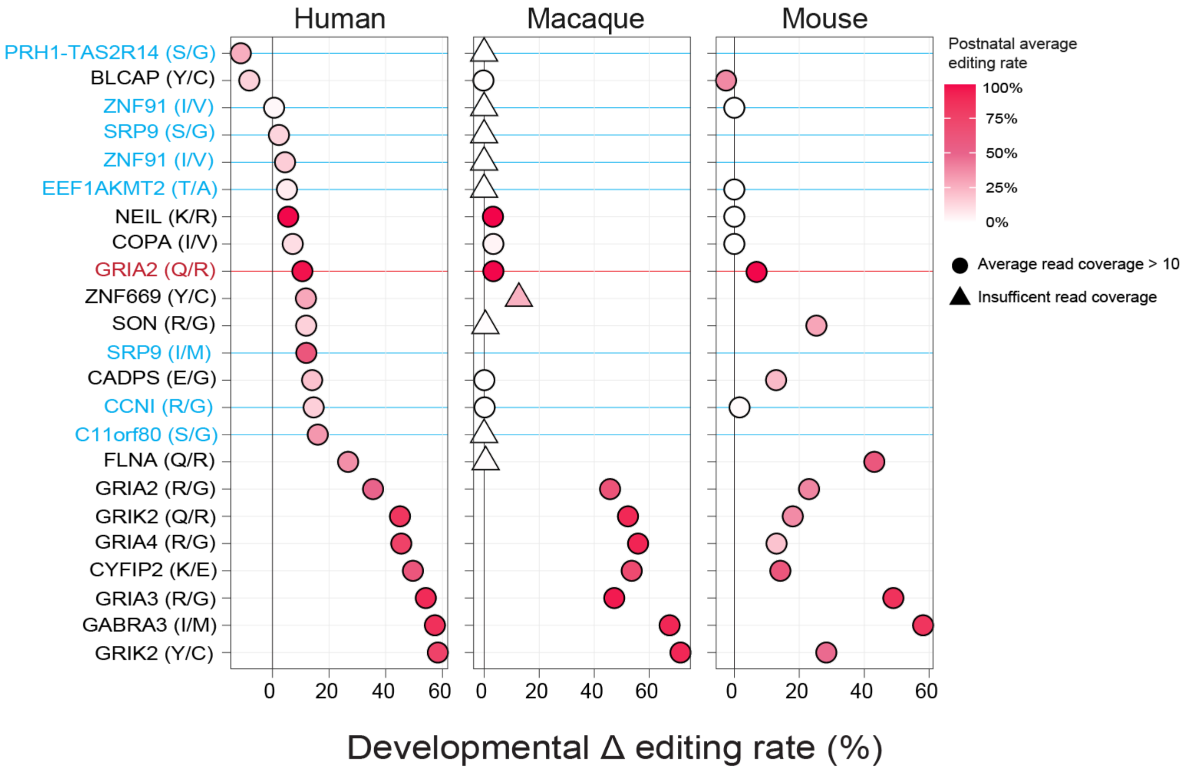


**Supplemental Figure 26. Developmental recoding sites across human, macaque and mouse.** We identified 23 known A-to-G recoding sites with significant developmental regulation in BrainVar (FDR P < 0.05) (left panel). Here we query these editing events across macaque (center panel) and mouse (right panel). The developmental change (delta) in editing rates is shown as a percentage, comparing the average editing rates from prenatal to postnatal periods. Each site is shaded by the average editing rate observed postnatally and shapes reflect whether that site had sufficient coverage across 70% of samples in the respective data set. Blue lines indicate potential human specific events. The red line denotes the classic Q/R site on *GRIA2*.

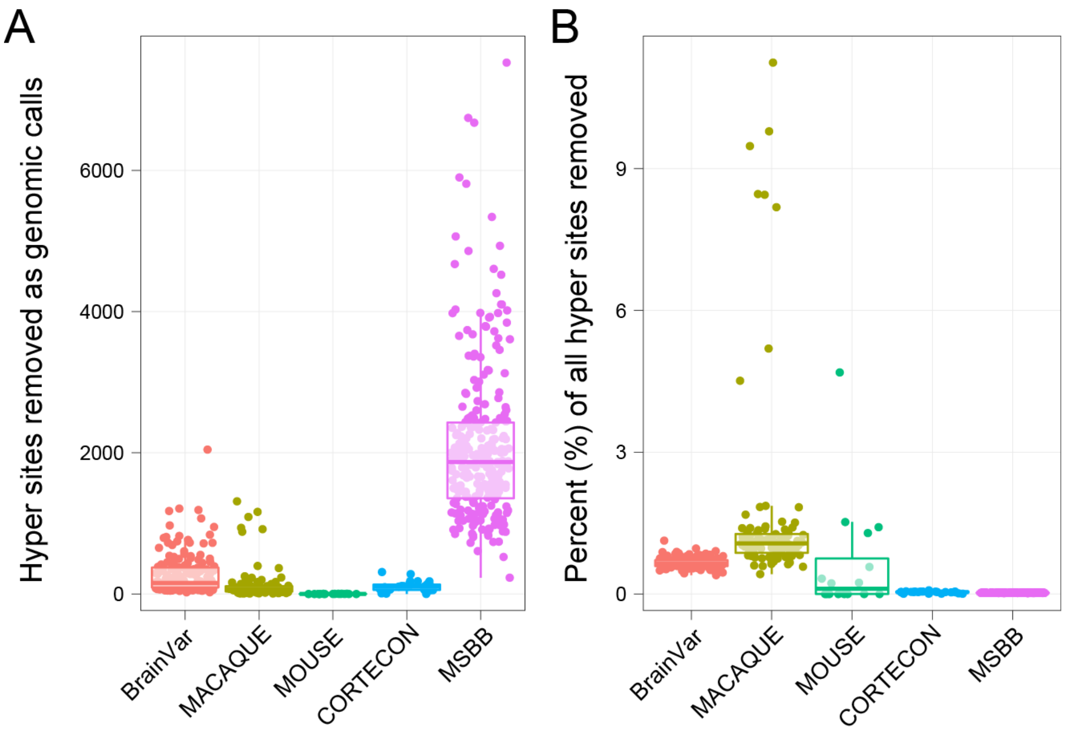


**Supplemental Figure 27. Number of genomic sites removed from the hyper-editing analysis. (A)** The total number of A-to-G hyper-editing sites and the (**B**) percentage (%) of all sites removed from the grant total (%) across all BrainVar, hESCs, MSBB, rhesus macaque and mouse samples. Notably, in addition to filtering for common genetic variants in dbSNP, BrainVar underwent additional filtering for private genomic variation using paired whole genome sequencing data. Overall, less than 2% of all hyper-editing sites were flagged and removed before proceeding to downstream analyses.

**Table S1. Alu editing index (AEI) measurements across all samples used in the current study.** This table also includes ADAR expression, predicted cell type proportions and corresponding metadata files used in the current study.

**Table S3. Differentially regulated RNA editing sites.** This table includes annotations and linear regression results for all BrainVar common sites, and their association with developmental age. We also provide the following information: 1) WGCNA module assignments; 2) WGCNA module enrichment and gene ontology (GO) reduction analysis; 3) module enrichment results for neuropsychiatric gene lists; 4) recoding site enrichment and GO reduction analysis.

**Table S4. Detection rates of differentially regulated RNA editing sites in independent data sets.** This table includes annotations as well as detection and coverage rates for all BrainVar common sites in MSBB and hESC common sites.

**Table S5. RNA hyper-editing results.** This table includes information on 1) raw unfiltered hyper-editing rates; 2) hyper-editing rates per sample in BrainVar, MSBB and CORTECON data sets, respectively; and 3) hyper-editing rate analysis of genes that accumulate hyper-editing sites over development.

**Table S6. RNA hyper-editing and splicing.** This table includes information on 1) the number of intronic RNA hyper-editing events per donor that are predicted to be splice alternating by SpliceAI and 2) information on intron retention rates per sample across development.

**Table S7. AEI and hyper-editing validation in animal models of neurodevelopment.** This table includes information on 1) AEI and hyper-editing rates in macaque cortical tissue and 2) AEI and hyper-editing rates in mouse whole cortex. Corresponding metadata are included.

**Table S8. Top temporal edQTL lists.** This table includes information for most significant editing-variant pair, covering 1) all constant edQTLs; 2) all prenatal predominate edQTLs and 3) all postnatal predominate edQTLs.

**Table S9. GWAS-edQTL co-localization results.** This table includes information on 1) the different PheCodes that were included from the UKBioBank; 2) a summary of co-localized hits based on constant, prenatal or postnatal predominate edQTLs; 3) a summary of all co-localized hits with PPH > 0.3. Results are separated for A-to-G edits relative to all other types of editing, which were treated as exploratory.

**References (specific to Supplemental Materials)**

References listed from number 51 to number 85 in the main text are unique to the Supplemental Material.
